# Structural basis for RNA-duplex unwinding by the DEAD-box helicase DbpA

**DOI:** 10.1101/2022.02.23.481582

**Authors:** Jan Philip Wurm

## Abstract

DEAD-box RNA helicases are implicated in most aspects of RNA biology, where these enzymes unwind short RNA duplexes in an ATP-dependent manner. During the central step of the unwinding cycle, the two domains of the helicase core form a distinct closed conformation that destabilizes the RNA duplex, which ultimately leads to duplex melting. Despite the importance of this step for the unwinding process no high resolution structures of this state are available. Here, we employ nuclear magnetic resonance spectroscopy and X-ray crystallography to determine structures of the DEAD- box helicase DbpA in the closed conformation, complexed with substrate duplexes and single- stranded unwinding product. These structures reveal that DbpA initiates duplex unwinding by interacting with up to three base-paired nucleotides and a 5’ single-stranded RNA duplex overhang. These high-resolution snapshots, together with biochemical assays, rationalize the destabilization of the RNA duplex and are integrated into a conclusive model of the unwinding process.

## Introduction

RNA molecules are involved in numerous cellular processes and their correct folding is often critical for function. RNA helicases that act as RNA chaperons, resolve misfolded RNA structures and rearrange RNA-protein complexes consequently being essential for cellular survival (Jarmoskaite and Russell, 2014).

The largest family of RNA helicases in eukaryotes is represented by DEAD-box proteins (Fairman- Williams et al., 2010). These enzymes unwind short RNA duplexes (up to 12-16 base pairs) in a non-processive and ATP-dependent manner. The functional core of DEAD-box helicases consists of two RecA like domains (termed RecA_N and RecA_C for the N- and C-terminal domains, respectively). The two domains are connected by a short, flexible linker and undergo large conformational changes during the unwinding cycle (Fig. 1a) (Linder and Jankowsky, 2011; Putnam and Jankowsky, 2013). In the apo-state, the helicase core adopts an open conformation, where the two RecA domains tumble independently (Sun et al., 2014; Theissen et al., 2008; Wurm, 2020) and where ATP can bind to the RecA_N domain (Mallam et al., 2012; Samatanga and Klostermeier, 2014). Binding of substrate RNA - in addition to ATP - induces the formation of a distinct closed conformation (Sun et al., 2014; Theissen et al., 2008), where ATP is sandwiched between the two domains and one of the RNA strands is bound to a bipartite active site, formed by both core domains (Sengoku et al., 2006). RNA binding to the active site leads to the destabilization of the RNA duplex and spontaneous dissociation of the unbound RNA strand (Rogers et al., 1999; Yang et al., 2007). The ATPase activity of DEAD-box helicases is greatly increased in the closed conformation. Upon ATP hydrolysis the helicase core returns to the open conformation (Theissen et al., 2008), while ADP and the single-stranded RNA product dissociate due to the dissolution of the active site. ATP binding is therefore sufficient for unwinding, with ATP hydrolysis mainly serving to release the helicase from unwound ssRNA (Liu et al., 2008). In addition to the productive unwinding cycle described above, DEAD-box helicases are prone to futile cycles, which occur when ATP is hydrolyzed prior to dissociation of the RNA duplex (Fig. 1a). These futile cycles increase sharply for longer, more stable duplexes due to decreased duplex dissociation rates (Putnam and Jankowsky, 2013).

**Fig. 1:**
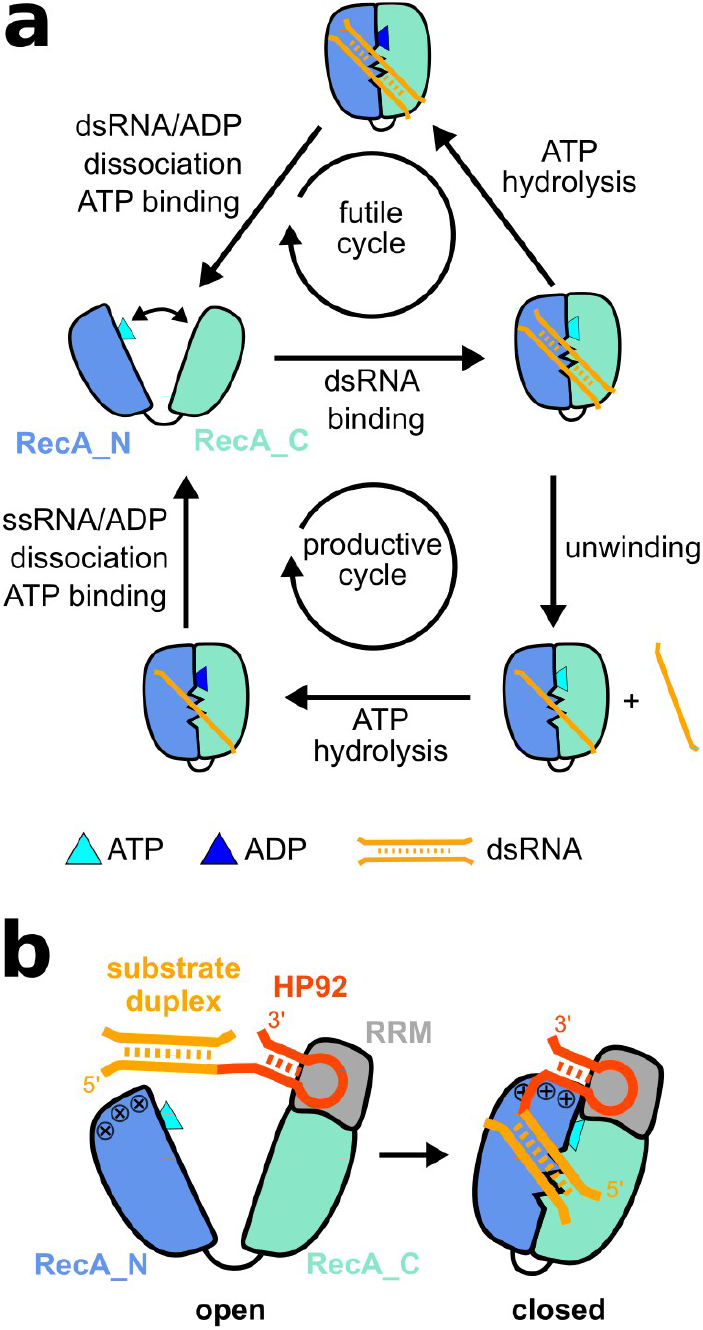
Unwinding cycle of DEAD-box helicases and activation of DbpA by HP92 RNA. (**a**) Schematic diagram of futile cycles (top) and productive unwinding cycles (bottom) of DEAD-box helicases. The helicase core (RecA_N domain blue, RecA_C domain cyan) alternates between an open conformation in the apo or ATP bound state and a closed conformation in the presence of ATP and RNA. (**b**) Domain orientation of DbpA in the open (left) and closed state (right). The C- terminal RRM (grey) orients HP92 (red) such that the stem of HP92 forms favorable interactions with a positively charged patch (indicated by + signs) on the RecA_N domain in the closed state. This stabilizes the closed state and enables the recruitment of the substrate duplex (orange, located 5’ to HP92) to the active site of the helicase core.

Several structures of DEAD-box helicases in the closed state, bound to single-stranded RNA (ssRNA) and non-hydrolyzable ATP analogs were determined in the last 15 years (Chen et al., 2020; Collins et al., 2009; Del Campo and Lambowitz, 2009; Montpetit et al., 2011; Ngo et al., 2019; Ren et al., 2017; Sengoku et al., 2006; Wong et al., 2016). Consequently, such structures correspond to the product-bound state following duplex unwinding and prior to ATP hydrolysis. The structures revealed a conserved ssRNA binding mode (Figure 1—figure supplement 1), where the bipartite, active site of the helicase core interacts with the sugar and phosphate backbone of six single- stranded nucleotides (nt). The RNA adopts a bent conformation due to the interaction between the RNA backbone and several highly conserved sequence motifs within the helicase core domains.

This RNA conformation strongly differs from the canonical A-form helical conformation and is thus incompatible with duplex formation (Russell et al., 2013). In agreement with the sequence- independent unwinding activity of DEAD-box helicases, no interactions are formed between the active site and RNA bases. In contrast to this wealth of structural information for the product-bound state, a structure of the closed-conformation complexed with double stranded RNA (dsRNA) substrate and ATP analog, is still missing. It is therefore unclear how the central step of the unwinding process - namely, the destabilization of the RNA duplex - is achieved at the molecular level.

A major challenge in the structural characterization of the destabilized duplex state lies in the trapping of this inherently unstable state in a homogeneous conformation. This is further complicated by the lack of sequence-specificity that facilitates binding in differing registers to the RNA. In order to overcome these challenges we selected the *E. coli* DEAD-box helicase DbpA for structural studies. DbpA possesses a C-terminal RNA recognition motif (RRM) (Wang et al., 2006) in addition to the helicase core and, we recently demonstrated that this RRM domain is stably anchored to the RecA_C domain (Fig. 1b) (Wurm et al., 2021). DbpA is involved in ribosome maturation (Sharpe Elles et al., 2009) and is recruited to the nascent ribosome by a specific, high- affinity interaction between the RRM and hairpin 92 (HP92) of the 23S rRNA (Fig. 1b) (Diges and Uhlenbeck, 2001; Hardin et al., 2010). Binding of HP92 is necessary for the helicase activity of DbpA, as a direct interaction between RRM-bound HP92 and the RecA_N domain strongly stabilizes - and thereby enables - formation of the closed conformation (Wurm et al., 2021). A substrate duplex linked to HP92 is therefore recruited to DbpA with high affinity and can also be efficiently unwound. Meanwhile, attachment onto HP92 locks the substrate duplex in a well-defined position relative to the active site of the helicase core consequently stabilizing the helicase/substrate complex.

In this study we employ nuclear magnetic resonance (NMR) spectroscopy to identify suitable ATP analogs and RNA constructs that enable trapping of DbpA in complex with a destabilized duplex. Based on these results we solved crystal structures of DbpA in the closed state, where the active site of the helicase core interacts with two different substrate RNAs: an RNA hairpin loop and a ss/dsRNA junction. In addition, we determined the DbpA structure bound to a ssRNA product. These structures can be readily integrated into a model of the unwinding cycle, where interactions between helicase and the ss/dsRNA junction rationalize how DbpA achieves the RNA duplex destabilization and promotes unwinding.

## Results

### A hairpin RNA substrate in combination with ADP/BeF_3_ traps the destabilized duplex state

In order to gain insights into the molecular mechanism that underlies duplex destabilization by DbpA during unwinding, we set out to identify conditions that trap the helicase when complexed with an unwinding intermediate. Since ATP binding is sufficient for unwinding, with ATP hydrolysis solely required for helicase recycling (Liu et al., 2008), we first screened for suitable non-hydrolyzable ATP analogs that support the formation of the closed state, based on their ability to fuel duplex unwinding by DbpA during single-turnover unwinding assays. To allow for unwinding by DbpA, the RNA construct included HP92 together with the substrate duplex consisting of nine bp (Fig. 2a). From the four investigated ATP analogs (ADPNP, ADPCP, ATPγS and ADP/BeF_3_), only ATPγS (k_obs_ 0.04 ± 0.01 min^-1^) and ADP/BeF_3_ (k_obs_ 0.10 ± 0.01 min^-1^) supported unwinding of the RNA substrate on a similar timescale as ATP (k_obs_ 0.43 ± 0.02 min^-1^). Interestingly, ATPγS was hydrolyzed during the unwinding process (Figure 2 - figure supplement 1) and is therefore not suitable for trapping the destabilized duplex over extended periods. Based on these results we identified ADP/BeF_3_ as a suitable ATP analogue for further investigations.

**Fig. 2:**
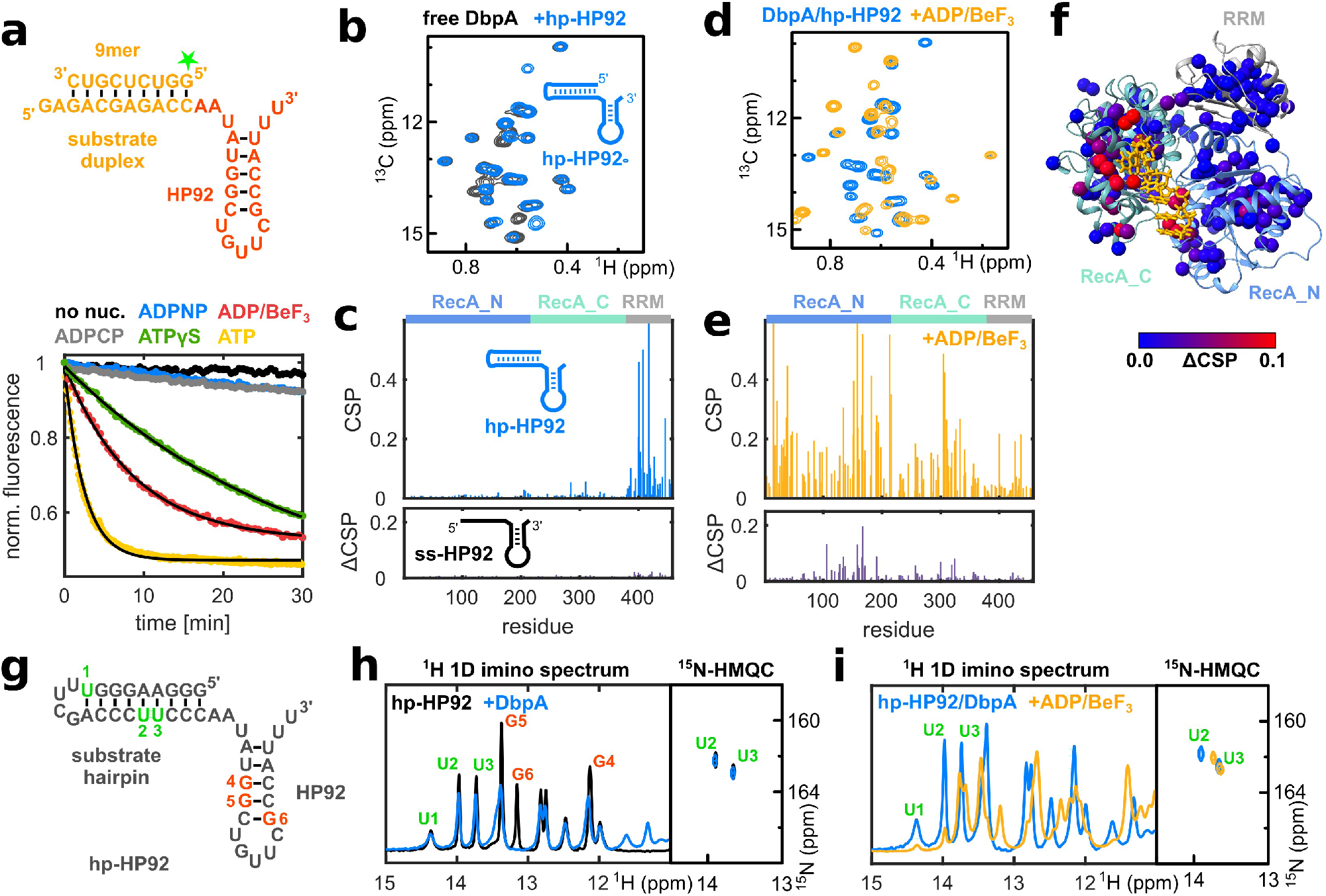
The DbpA/hp-HP92/ADPBeF_3_ complex represents a trapped unwinding intermediate. (**a**) Fluorescence based unwinding assays in the presence of different ATP analogs. A 5’ fluorescein (green star) labeled 9mer RNA is hybridized to an RNA containing HP92 (top). Unwinding can be followed by a decrease in fluorescence intensity (bottom). Mono-exponential fits to the fluorescence time traces are shown in black for ATP, ADP/BeF_3_ and ATPγS. (**b**) Ile region of methyl TROSY spectra of ILMVA-labeled DbpA in the free state (black) and bound to hp-HP92 RNA (blue). (**c**) Sequence plots of CSPs induced by binding of the hp-HP92 RNA (top) and differences between CSPs (ΔCSPs) induced by binding of the ss-HP92 RNA and the hp-HP92 (bottom). DbpA domains are indicated at the top. (**d**) Methyl TROSY spectra of DbpA/hp-HP92 complex prior to (blue) and after addition of ADP/BeF_3_ (orange). (**e**) Sequence plots of CSPs induced by binding of ADP/BeF_3_ to the DbpA/hp-HP92 RNA complex (top) and ΔCSPs between binding of ADP/BeF_3_ to the ss-HP92/DbpA and to the hp-HP92/DbpA complexes (bottom). (**f**) ΔCSPs from (e) plotted onto a model of the closed state of DbpA bound to ssRNA ((Wurm et al., 2021), red large ΔCSPs, blue small ΔCSPs). The ssRNA is shown in orange. (**g**) Sequence of hp-HP92 RNA. Nucleotides with assigned imino proton signals are numbered and shown in red (HP92) or green (substrate hairpin). (**h**) ^1^H-1D imino proton spectra (left) and ^15^N-HMQC spectra (right) of uridine ^15^N labeled hp-HP92 RNA prior to (black) and after addition of DbpA (blue). (**i**) Same spectra as in (h), but for the hp- HP92/DbpA complex prior to (blue) and after addition of ADP/BeF_3_ (orange).

To enforce trapping of the destabilized duplex by DbpA and to prevent the dissociation of the RNA duplex, we made use of an RNA construct (hp-HP92 RNA) containing a hairpin structure with a stable UUCG tetraloop as the substrate (Fig. 2B). This substrate hairpin prevents the dissociation of the destabilized duplex. To probe formation of the closed state, we performed NMR titrations using deuterated DbpA that is ^1^H,^13^C-labeled at the methyl-groups of Ile, Leu, Met, Val, Ala (ILMVA) and that yields high-quality methyl-group spectra ((Wurm et al., 2021); Figure 2 - figure supplement 2).

Binding of the hp-HP92 RNA substrate to DbpA can be monitored based on changes (chemical shift perturbations CSPs) of the position of the methyl-group signals (Fig. 2b, Figure 2 - figure supplement 2). In the absence of ATP, RNA binding induces large CSPs for residues that belong to the RRM, though only minimally affecting signals from the RecA core domains (Fig. 2c, upper graph). This demonstrates that only the RRM of DbpA interacts with HP92 of the hp-HP92 RNA, and that no specific contacts between the substrate duplex and the core domains are formed in the absence of ATP analog. In line with this finding, essentially identical CSPs are elicited by an RNA construct (termed ss-HP92) that contains a single-stranded region instead of the substrate duplex (Fig. 2c, bottom graph, note that differences in CSPs (ΔCSPs) are shown, Figure 2 - figure supplement 3).

During the next step we added the identified ATP analogue ADP/BeF_3_ to the DbpA/hp-HP92 complex. This leads to large CSPs within the helicase core domains (Fig. 2d, 2e top graph, Figure 2 - figure supplement 4). As we have shown previously (Wurm et al., 2021) these CSPs mark the formation of the DbpA closed state. Similarly, the closed state is also formed upon addition of ADP/BeF_3_ to the DbpA/ss-HP92 complex (Figure 2 - figure supplement 5). Interestingly, the DbpA spectra for the closed state reveal several distinct differences between ss-HP92 and hp-HP92 RNA complexes (Fig. 2e, bottom graph, Figure 2 - figure supplement 5) and these differences cluster in the vicinity of the helicase core active site (Fig. 2f). We reasoned that these differences arise from the interaction with the intact substrate duplex in the hp-HP92 complex rather than the single- stranded region in the ss-HP92 RNA suggesting that the substrate duplex is not completely unwound in the hp-HP92/DbpA complex.

To verify the presence of the substrate duplex in the closed state we turned to 1D ^1^H imino proton spectra (Fig. 2g,h). Only imino protons of nucleotides that are involved in stable base pairs give rise to observable signals in such spectra. To confirm the presence of the substrate hairpin we concentrated on the three uridines within this hairpin (labeled U1 to U3 in Fig. 2g,h). Binding of DbpA in the absence ADP/BeF_3_ only leads to a slight reduction in signal intensity, due to the increased molecular weight of the complex. This shows that there is no (or only a highly transient) interaction between DbpA and the substrate hairpin of the hp-HP92 RNA. By contrast, the guanosine residues of the HP92 stem (labeled G4-G6 in Fig. 2g,h) show clear CSPs due to direct interaction with the RRM of DbpA (Hardin et al., 2010; Wurm et al., 2021). Addition of ADP/BeF_3_ to the DbpA/RNA complex leads to formation of the closed state of DbpA and induces large shifts for essentially all imino proton signals (Fig. 2i). This provides clear evidence for a direct interaction between DbpA and the substrate hairpin within the closed state. At least 7-8 of the nine imino proton signals are still visible indicating that the substrate hairpin is still largely intact when bound to the active site of DbpA. This is supported by ^1^H^15^N-HMQC spectra of uridine ^15^N-labeled hp- HP92 RNA (Fig. 2i). In these spectra only imino signals of based paired uridines are detectable. The spectra show two uridine imino signals in the closed state, which must originate from the substrate duplex as we have shown previously that the uridine imino signals of HP92 are not detectable under this measurement conditions (Wurm et al., 2021). We tentatively assign the signals to the central uridines of the substrate hairpin (U2/U3).

In summary, these results demonstrate that the DbpA/hp-HP92/ADP/BeF_3_ complex represents a trapped unwinding intermediate where the mostly-intact duplex of the substrate hairpin interacts with the active site of the helicase core.

### Structure of the hairpin RNA bound to the active site of DbpA

To understand how duplex destabilization by DbpA is achieved on a structural level, we solved the crystal structure of the DbpA/hp-HP92/ADP/BeF_3_ complex to a resolution of 3.2 A (Fig. 3a, Figure 3 - figure supplement 1). Surprisingly, a 2:2 DbpA:RNA complex is formed in the crystal. Each DbpA molecule interacts with the HP92 of one RNA molecule (via the RRM) and with the substrate hairpin (via the core domains’ active site) of the other RNA. The two hairpins of each RNA molecule stack co-axially and HP92 is elongated by 2 bp, which are formed between the 3’ overhang of HP92 and the linker between HP92 and the substrate hairpin (nt 23-25, see Figure 3 - figure supplement 2 for details). Since the good quality of the NMR spectra (Fig. 2d) strongly argued against a 2:2 complex with a molecular weight >120 kDa, we investigated the oligomeric state of the DbpA/hp-HP92/ADP/BeF_3_ complex in solution, using size exclusion chromatography (SEC) experiments (Fig 3c). The experiments show that addition of ADP/BeF_3_ to the DbpA/hp- HP92 complex leads to the formation of two species: A minor population of a larger species, which would be compatible with a 2:2 complex and a major population of a more compact species, which we assign to the 1:1 complex. These results indicate that 1:1 and 2:2 complexes co-exist in solution. We conclude that our NMR experiments predominantly report on the major population of the 1:1 complex, whereas the minor population of the 2:2 complex was crystallized. Notably, the SEC and NMR experiments were conducted at similar concentrations, with higher concentrations used for crystallization trials, which is expected to increase the population of the 2:2 complex. Importantly, molecular modeling establishes that a 1:1 complex with identical DbpA/RNA interactions as observed for the 2:2 complex can be formed (Fig. 3b). We are thus confident that the observed interactions in the crystal structure between DbpA and the RNA hairpins are relevant in solution.

**Fig. 3:**
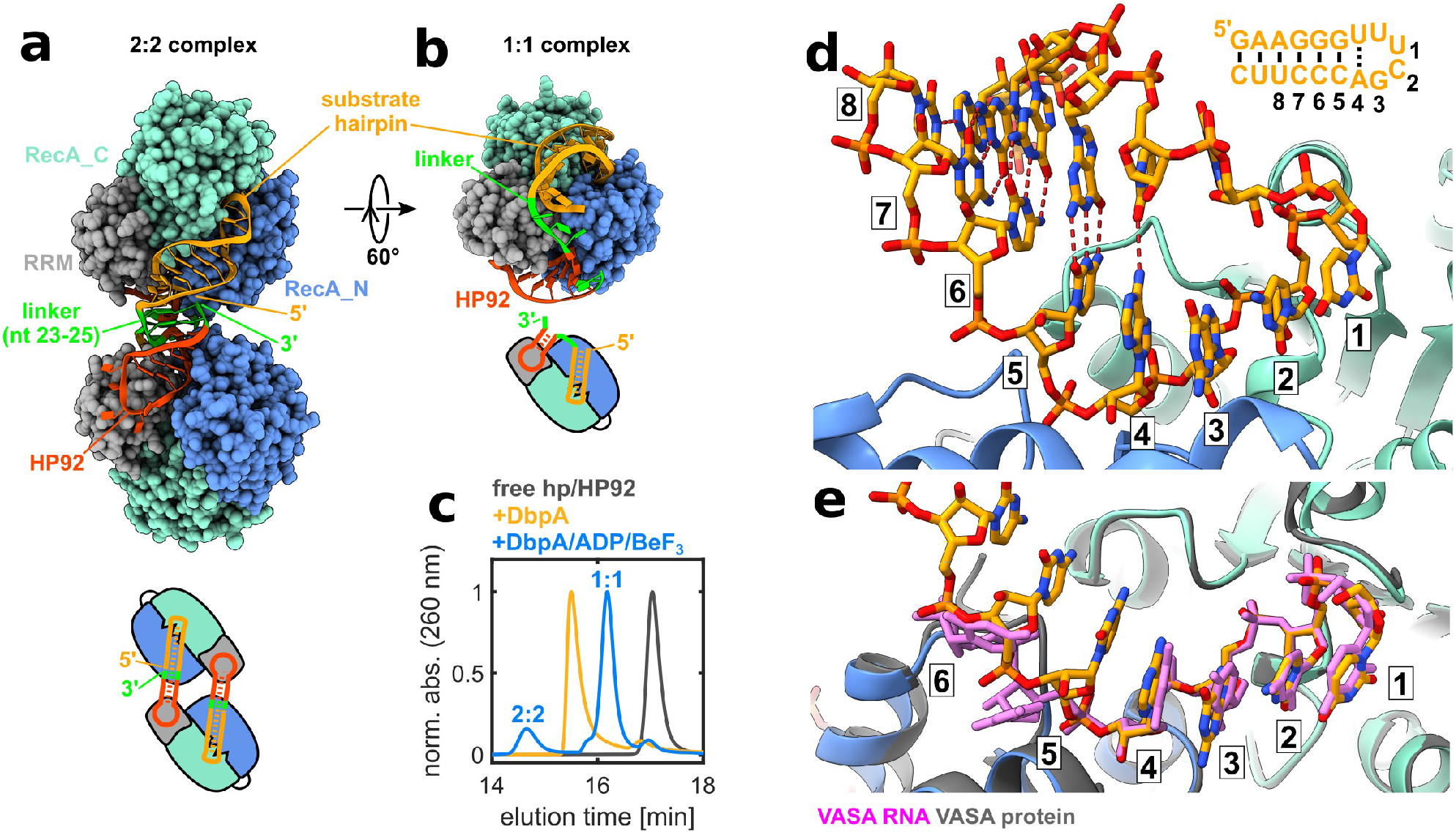
Structure of the hp-HP92/DbpA complex in the closed state. (**a**) Overall structure (top) and schematic diagram (bottom) of the complex between two DbpA molecules (chains A/B) and two hp-HP92 RNAs (chains F/G), as observed in the crystal. HP92 (red), the substrate hairpin (orange) and the 3 nt linker that base pairs with the 3’ overhang of HP92 (green) are shown in ribbon representation. The RecA_N (blue), RecA_C (cyan) and RRM (grey) domains of DbpA are shown in sphere representation. (**b**) Model of the 1:1 complex obtained by connecting HP92 and the substrate hairpin bound to one DbpA molecule by a flexible 3 nt linker (green). (**c**) SEC chromatograms of free hp-HP92 (black), the hp-HP92/DbpA complex (orange) and the hp-HP92/DbpA/ADP/BeF_3_ (blue) complex. (**d**) Close-up of the interaction between substrate hairpin (chain F, orange) and the active site of the helicase core (chain B, RecA_N blue, RecA_C cyan). The nucleotides that interact with the active site are numbered 1-6. The substrate hairpin is shown in the upper-right, with identical numbering. The distorted base pair between the adenosine in position 4 and the opposite uridine is indicated by a dashed line. (**e**) Comparison between the substrate hairpin bound to DbpA (only nt 1-7 are shown for clarity) and the complex between a 6 nt ssRNA (pink) and the DEAD-box helicase VASA (grey) (PDB ID 2db3).

The interactions between HP92 and the RRM of DbpA are basically identical to the interactions observed for a homologous RRM/HP92 complex that was previously published (Hardin et al., 2010) (Figure 3 - figure supplement 3) and we hereby focus on the interaction between the substrate hairpin and the active site of the helicase core (Fig. 3d, Figure 3 - figure supplement 4a). The split active site interacts with 6 nt (numbered 1-6 in Fig. 3d) of the hairpin. The first 3 nt belong to the loop of the substrate hairpin, whereas nt 4-6 are part of the stem. Due to interaction with the active site, the first base pair of the stem (formed between the adenosine at position 4 and the opposing uridine) is distorted in comparison to a regular AU base pair, and consequently only forms one hydrogen bond. Conversely, the cytosines in position 5 and 6 form regular GC base pairs with the opposing strand. All six bases in the binding site show continuous base stacking interactions.

Next, we compared the RNA conformation within the hp-HP92/DbpA complex with the canonical ssRNA product-bound state of other DEAD-box helicases (exemplified by the VASA helicase (Sengoku et al., 2006)) The conformation of nt 1-4 in both structures is basically identical. The main differences reside in the conformation of nt 5 and 6. In the VASA structure the bases of nt 5 and 6 are rotated by 90° relative to the DbpA structure and consequently their conformation is not compatible with duplex formation. In contrast to the known, product-bound states, our structure thus demonstrates that DbpA is able to bind a ss/dsRNA junction that contains a 5’ ssRNA overhang of 3 nt, whereby the 3 nt overhang and the first 3 nt of the duplex interact with positions 1-3 and 4-6 of the active site. We propose that the crystallized conformation represents a trapped unwinding intermediate that captures the interaction between a ds/ssRNA junction and DbpA during unwinding.

### Structure of a ss/dsRNA junction bound to the active site of DbpA

To exclude the possibility that the observed RNA conformation is influenced by the short loop of the substrate hairpin, we crystallized DbpA with a permuted RNA construct (termed ds-HP92) that contains a ss/dsRNA junction with a 3 nt 5’ overhang instead of the hairpin. To this end, the two co- axially stacking helices of the hp-HP92 RNA were fused, which places the 5’ and 3’ ends at the position of the loop of the substrate hairpin (Fig 4a, Figure 3 - figure supplement 2c). The complex crystallized as a 2:2 complex similar to the hp-HP92 RNA (Fig 4a, Figure 3 - figure supplement 2d) and the structure was determined to a resolution of 3.0 Å (Figure 3 - figure supplement 1). Since fusion of the helices prevents the substrate duplex from folding back onto the active site of the same DbpA molecule, the 2:2 complex is also the major species in solution (Fig. 4b). In the structure, the 3 nt 5’ overhang and the first 3 nt of the substrate duplex are bound to the active site in positions 1-3 and 4-6, respectively (Fig. 4c, Figure 3 - figure supplement 4b). The DbpA/RNA interactions are virtually identical to the ones observed in the hairpin structure, with the exception that the first base pair of the duplex (formed between the adenosine at position 4 and the opposing uridine) is not distorted. The distortion in the first base pair of the hp-HP92 RNA thus most likely originates from the strain in the short loop of the substrate hairpin.

**Fig. 4:**
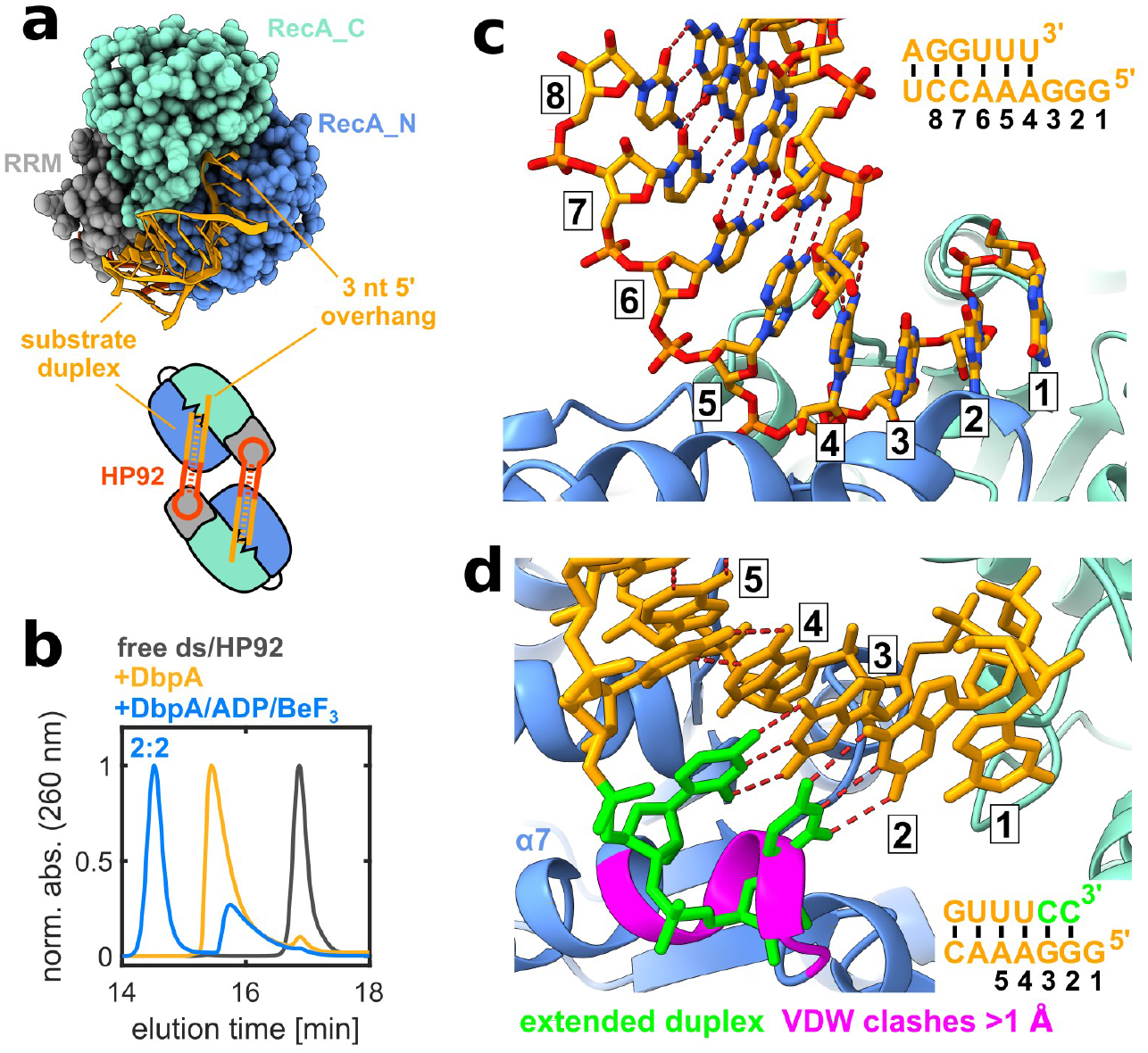
Structure of the ds-HP92/DbpA complex in the closed state. (**a**) Interaction of the substrate duplex including a 3 nt 5’ overhang with the active site of the helicase core. For clarity, only the substrate duplex (chain D) and one DbpA molecule (chain B) are shown (top). Schematic diagram of the 2:2 complex observed in the crystal (bottom). (**b**) SEC chromatograms of free ds- HP92 (black), the ds-HP92/DbpA complex (orange) and the ds-HP92/DbpA/ADP/BeF_3_ (blue) complex. (**c**) Close-up of the interaction between the substrate duplex (chain D, orange) and the active site of the helicase core (chain B, RecA_N blue, RecA_C cyan). The nucleotides that interact with the active site are numbered 1-6. The substrate duplex is shown in the upper-right, with identical numbering. (**d**) Extension of the duplex by addition of 2 nt at the 3’ end (green) leads to severe clashes with α-helix α7 of the RecA_N domain. Residues that show van der Waals clashes > 1 Å with the RNA are shown in pink.

To evaluate whether an extended duplex with base-paired nucleotides in positions 2 and 3 could interact with DbpA in a similar manner, we modeled two additional nucleotides at the 3’ end of the RNA in an A-form helix conformation (Fig. 4d, green residues). The model reveals that both nucleotides of the extended helix would severely clash with α-helix α7 of the RecA_N domain. This indicates that: i) a 5’ single-stranded region of 3 nt is necessary for the interaction of duplex RNA with DbpA; ii) only nucleotides in positions 4-6 can be part of a duplex, whereas positions 1-3 are not compatible with dsRNA. In summary, these results enforce the notion that DbpA interacts with a ds/ssRNA junction during the unwinding process, where nucleotides in position 1-3 are single- stranded and nucleotides 4-6 can be part of a duplex.

### Structure of the ssRNA product bound to the active site of DbpA

To complete the picture of the unwinding cycle of DbpA, we aimed to also gain insights into the ssRNA product-bound state after duplex unwinding. To this end we solved the structure of DbpA in complex with ADP/BeF_3_ and an RNA construct termed ss-HP92 (Figure 3 - figure supplement 2e). This RNA contains a single-stranded region 5’ to HP92 that interacts with the active site of the helicase core (Fig. 5a). Again crystallization resulted in a 2:2 RNA:DbpA complex (Figure 3 - figure supplement 2f) and SEC experiments show that the 2:2 complex is predominant in solution (Fig. 5b). Similar to the ds-HP92 and hp-HP92 complexes, each DbpA molecule interacts with HP92 of one RNA via the RRM and with the single-stranded region of the other RNA via the core domains. Three 2:2 complexes are present in the asymmetric unit. All of them display highly similar structures. The main differences reside in the conformation of the single-stranded region of the RNA. Three RNAs (chains G, H and L) adopt the same canonical ssRNA binding mode as observed in other DEAD box helicases (conformation 1, Fig. 5C, Figure 3 - figure supplement 4c). The bases of nt 1-4 show a continuous stacking interaction, whereas the bases of nt 5 and 6 are rotated by 90° relative to nt 1-4. In the other three RNAs (chains C, D and K) all six bases show a continuous stacking interaction (conformation 2, Fig. 5d, Figure 3 - figure supplement 4d). This continuous base stacking and the overall RNA conformation resembles the ds-HP92/DbpA complex structure (Figure 5 - figure supplement 1). Conformation 2 could therefore represent a snapshot of the RNA conformation immediately following duplex dissociation, before the RNA transitions into conformation 1.

**Fig. 5:**
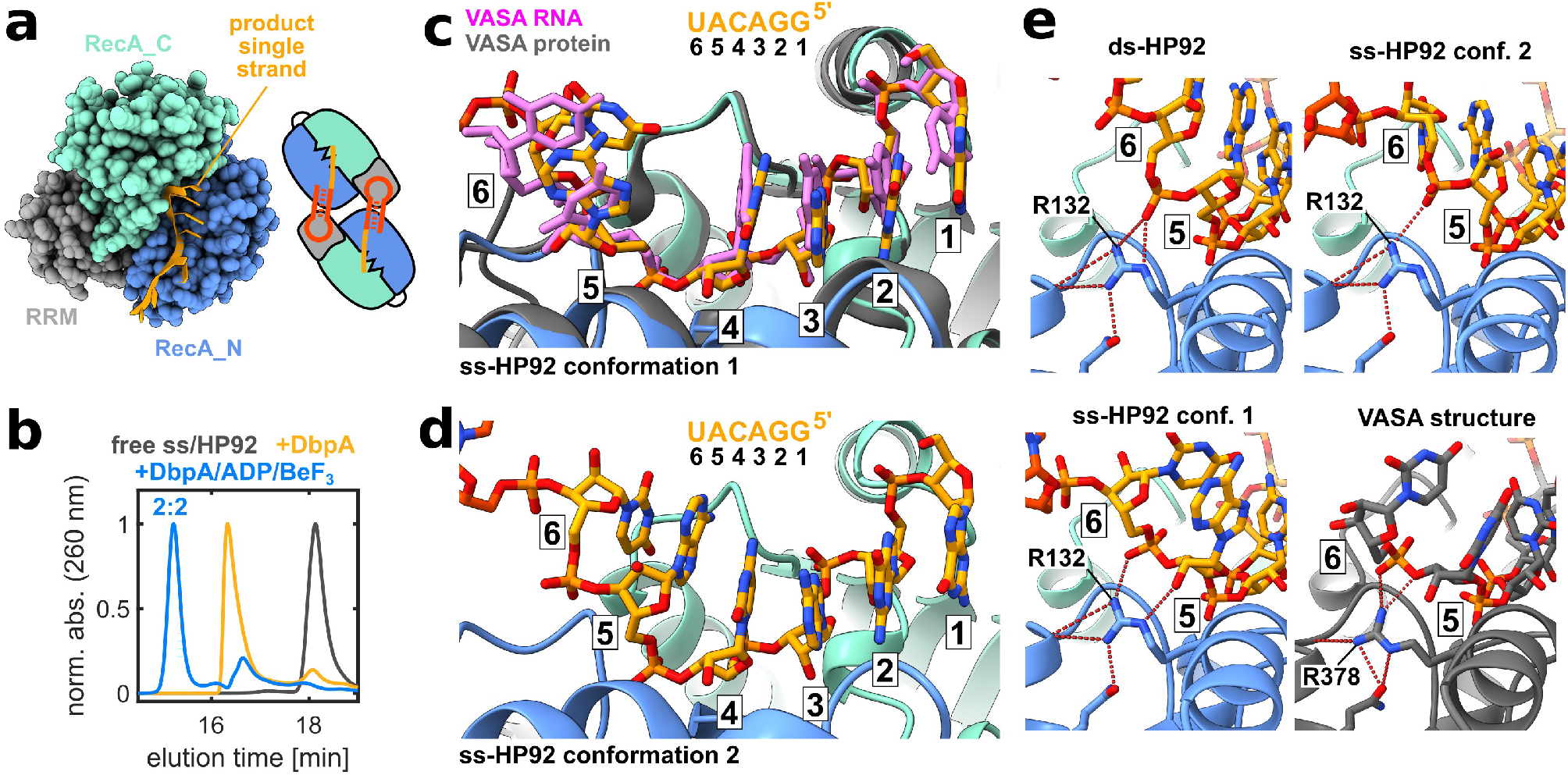
Structure of the ss-HP92/DbpA complex in the closed state. (**a**) Interaction of the ssRNA with the active site of the helicase core. For clarity, only the ssRNA (chain L) and one DbpA molecule (chain I) are shown (left). Schematic diagram of the 2:2 complex observed in the crystal (right). (**b**) SEC chromatograms of free ss-HP92 (black), the ss-HP92/DbpA complex (orange) and the ss-HP92/DbpA/ADP/BeF_3_ complex (blue). (**c**) Close-up of the interaction between the ssRNA region (chain L, orange) in conformation 1 and the active site of the helicase core (chain I, RecA_N blue, RecA_C cyan). For comparison, the complex between a 6 nt RNA (pink) and the DEAD-box helicase VASA (grey) (PDB ID 2db3) is shown. The nucleotides that interact with the active site are numbered 1-6. The ssRNA sequence is shown on top with identical numbering. (**d**) Close up of the interaction between the ssRNA region (chain C, orange) in conformation 2 and the active site of the helicase core (chain A, RecA_N blue, RecA_C cyan). (**e**) Comparison of the interactions between nucleotides in position 5 and 6 for DbpA and VASA. The top row and the bottom-left panel show complexes between DbpA and ds-HP92 and ss-HP92 in conformation 1 and 2. The lower-right panel shows the VASA/ssRNA complex (PDB ID 2db3). Hydrogen bonds formed by R132 (DbpA) or the corresponding R378 (VASA) are indicated by dashed-red lines.

These results demonstrate that DbpA conforms to the canonical ssRNA product binding mode observed for other DEAD-box helicases, but also suggest a notable plasticity in its ssRNA interaction mode. A comparison between the ds-HP92 and ss-HP92 RNA bound structures and the VASA/RNA complex shows that the interaction with nt 1-4 is essentially identical in all structures (Figure 5 - figure supplement 2) and that the differences reside mainly in the interaction of nt 5 and 6 with the helicase core (Fig. 5e). In the canonical binding mode, the conserved R378 of VASA forms hydrogen bonds with the ribose of nt 5 and the phosphate group of nt 6. In the DbpA structures, the corresponding R132 is slightly repositioned, but forms similar interactions with nt 5 and 6. We conclude that subtle structural differences in the conformation of R132 allow DbpA to favorably interact with duplex as well as with ssRNA at positions 4-6.

### Activity assays support an initial interaction with a ss/dsRNA junction during unwinding

In summary, the structures of the ds-HP92 and ss-HP92 complexes suggest the following model of duplex unwinding from a 5’ overhang. DbpA initially binds to the ss/dsRNA junction of the duplex, where 3 nt of the 5’ overhang interact with position 1-3 and the first 3 nt of the duplex with position 4-6. Breathing of the duplex ends, which takes place on the ms timescale (Snoussi and Leroy, 2001), thus leads to ss-HP92 conformation 2, and subsequently to ss-HP92 conformation 1. The ss- HP92 conformation 1 prevents duplex re-formation, thereby destabilizing the remaining duplex and accelerating its spontaneous dissociation.

To support this model, we performed single turnover unwinding assays with RNA constructs that contain a 9-bp duplex and a 5’ ssRNA overhang of 0, 2, 5 or 8 nt (Fig. 6a). The observed duplex unwinding rates (Fig. 6b) increase for overhangs of 0 and 2 nt from 0.20 min^-1^ to 0.42 min^-1^ and then decline for longer 5’ overhangs of 5 (0.32 min^-1^) and 8 nt (0.21 min^-1^). These results can be rationalized based on our model. The 5’ overhang of 0 nt shows the lowest unwinding rate, as formation of the closed state depends on the presence of a 5’ single-stranded region. In the absence of a 5’ ssRNA overhang, this single-stranded region is only transiently formed by fraying of the terminal base pairs. Elongation of the overhang to 2 nt increases unwinding, as the overhang facilitates the formation of the closed state and only fraying of the terminal base pair is necessary. Longer overhangs are expected to facilitate the formation of the closed state even further, but also increase the risk of unproductive interactions, where DbpA would only bind to the ssRNA overhang, which would not destabilize the duplex region.

**Fig. 6:**
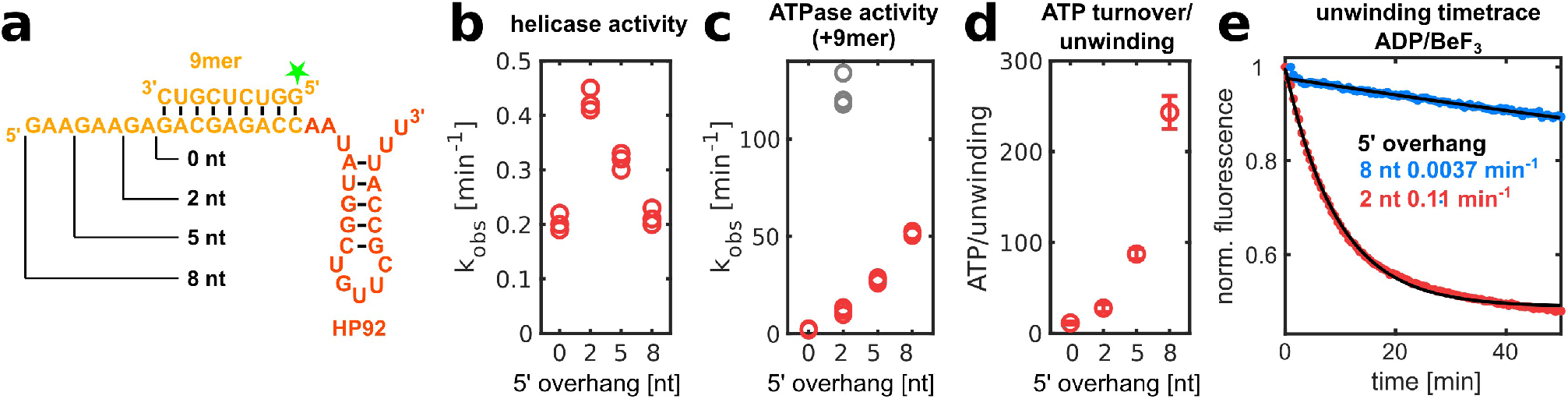
The length of the ssRNA 5’ overhang influences helicase and ATPase activity of DbpA. (**a**) RNA constructs used for activity assays. The substrate duplex (orange) is located 5’ to HP92 (red). The varying length of the 5’ overhangs is indicated. For helicase assays, the 9mer RNA contained a fluorescein label at the 5’ end (indicated by a green star). (**b**) Unwinding rates observed in single-turnover experiments are plotted versus the 5’ overhang length. Results from three measurements are shown. (**c**) ATPase turnover rates are plotted versus the 5’ overhang length. Rates were determined in the absence (grey) and presence of the 9mer RNA (red). Results from three measurements are shown. (**d**) The number of hydrolyzed ATP molecules for each unwinding event are plotted versus the 5’ overhang length. The values represent mean and standard deviation calculated from the experiments shown in panels b and c. (**e**) Single-turnover unwinding of RNA constructs with a 5’ overhang of two (red) or eight nt (blue) in the presence of ADP/BeF_3_ is followed by fluorescence intensity measurements. Exponential fits to fluorescence time traces are shown in black and the unwinding rates obtained from the fits are given.

This finding is corroborated by ATPase assays with identical RNA constructs including the 9-bp duplex (Fig. 6c, red symbols). ATP hydrolysis requires formation of the closed state and ATPase rates therefore report on the efficiency of closed state formation. The observed ATPase rates increase proportionally with 5’ overhang length from 2.3 min^-1^ for the 0 nt overhang to 52 min^-1^ for the 8 nt overhang, confirming that longer overhangs facilitate the formation of the closed state.

Consequently, constructs with longer 5’ overhangs require additional ATP hydrolysis events for unwinding (Fig. 6d) as DbpA mainly forms unproductive closed states, where it binds to the 5’ ssRNA overhang remote from the RNA duplex. Based on this reasoning, ADP/BeF_3_ should not support unwinding of duplexes with long 5’ overhangs, since DbpA should be trapped predominantly within unproductive closed states (note that a 5’ overhang of 2 nt was used in the helicase assays in Fig. 2a). This is experimentally verified in unwinding assays for the 8 nt 5’ overhang, where unwinding is strongly reduced compared to the 2 nt overhang (Fig. 6e). These results and the increased ATPase rate for longer 5’ overhangs indicate that DbpA does not favor the interaction with the ss/dsRNA junction over the interaction with the ssRNA 5’ overhang. To test this, we performed ATPase assays in the absence of the 9mer RNA, for the RNA carrying the 2 nt 5’ overhang (Fig. 6c, grey symbols). This leads to a 10.4-fold increase in the ATPase rate and indicates that formation of the closed state is even more favorable for a ssRNA substrate, compared to a ss/dsRNA junction. In summary, the observed helicase and ATPase rates are in good agreement with a model whereby DbpA preferentially forms the closed state upon interaction with a 5’ ssRNA overhang or a ssRNA/duplex junction, as observed in structures of the ss-HP92 and dsHP92 complexes.

### Transient active site formation and substrate RNA binding in the absence of ATP

Finally, we sought to gain direct insights into the initiation of the unwinding process, namely, the formation of the closed state. Closing of the DbpA/RNA complex is inherently inefficient as three bodies (the two RecA domains and substrate RNA) require assembly in the correct orientation. This raises the question whether the substrate RNA binds to a pre-formed active site (where the two RecA domains assemble initially) or whether the substrate RNA initially interacts with one of the RecA domains prior to the active site assembly. To test for active site pre-formation we performed paramagnetic relaxation enhancement (PRE) experiments using RNA that corresponds to isolated HP92 (Fig. 7a, Figure 7 - figure supplement 1a) and that carries a nitroxide spin label at the 3’ end. The spin label induces a PRE that leads to a strong decrease of the methyl-group NMR signals in its vicinity, which can be readily quantified and allow for the detection of sparsely populated states (Clore et al., 2007). Upon binding of the spin labeled HP92, we observe the expected PREs close to the HP92 binding site on the RRM. Additional PREs are observed on top of the RecA_N domain (Fig. 7a, Figure 7 - figure supplement 1a) indicating that this region of the RecA_N domain also approaches the spin label. Since close contacts between the RecA_N domain and HP92 are only found in the closed state of the enzyme (Fig. 7a), these results clearly indicate that DbpA transiently adopts the closed state in the presence of HP92. It would be interesting to explore whether the closed state is also formed in the absence of HP92. However, due to the nine native cysteine residues in DbpA we were not able to selectively introduce a spin label in the protein.

**Fig. 7:**
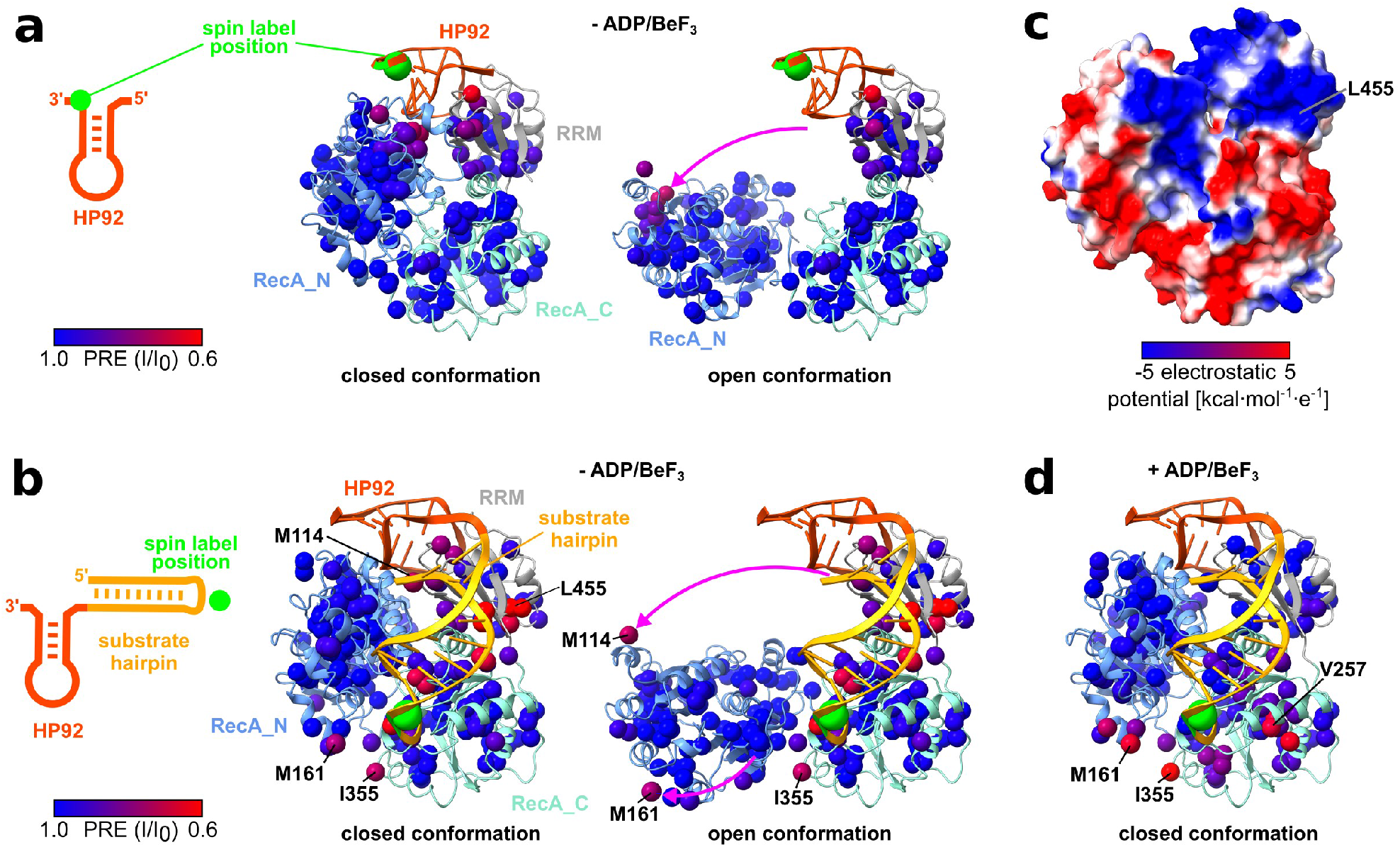
DbpA transiently samples the closed state in the absence of ATP. (**a**) PRE experiments were performed with spin labeled HP92 RNA and ILMVA-labeled DbpA in the absence of ADP/BeF_3_. The RNA construct is shown on the left. The position of the 4-thiouridine residue that carries the nitroxide spin label is indicated by a green circle. The methyl-groups of DbpA are colored according to the decrease in signal intensity, due to spatial proximity of the spin label from blue (no effect) to red (strong reduction). The closed conformation (left structure) and an arbitrary open conformation (right structure) are depicted. The pink arrow indicates the re-orientation of the RecA_N domain between the two structures. The C1’ atom of the spin labeled 4-thiouridine is shown as a green sphere. (**b**) PRE experiments with an hp-HP92 RNA containing a spin label in the loop of the substrate hairpin (green circle) (left). Methyl-groups are colored as in (a) for the closed- conformation (left structure) and for an arbitrary open conformation (right structure). The pink arrows indicates the re-orientation of the RecA_N domain. The methyl-groups of M114 and M161 that exhibit the largest PREs in the RecA_N domain are labeled. The RNA is shown in the conformation observed in the DbpA/hp-HP92 complex, where the substrate interacts with the active site of the helicase core. (**c**) DbpA in the closed conformation colored according to electrostatic surface potential (blue = negative, red = positive). (**d**) PRE experiments with identical RNA as in (b), but in the presence of ADP/BeF_3_. Methyl-groups are colored as in (a) and DbpA is shown in the closed conformation.

To test for interactions between the substrate duplex and the RecA domains, we repeated the PRE experiments with a hp-HP92 RNA construct that carries a spin label in the loop of the substrate hairpin (Fig. 7b, Figure 7 - figure supplement 1b). In the absence of ATP analog, the PREs for this RNA construct are distributed over a large surface on the RRM and RecA_C domains, indicating that the substrate hairpin only transiently interacts with these domains, and samples a large set of different conformations. The largest PREs are observed for a positively charged region on the RRM surrounding L455 (Fig. 7c). This positively charged patch is located remotely from the active site, and its function is currently unknown. It was speculated to be important for RNA interactions in the context of the pre-ribosome (Hardin et al., 2010) and most likely attracts the substrate hairpin in the PRE experiments due to its positive charge. More importantly, the residues from the RecA_N domain with the largest PREs (M114 and M161) and the residues with large PREs on the RecA_C domain are in close contact solely in the closed conformation (Fig 7b). This indicates that DbpA also transiently adopts the closed conformation in the presence of hp-HP92 and thereby corroborates our results for the isolated HP92 (Fig. 7a). In addition to M161 from the RecA_N domain, large PREs are also found for I355 from the RecA_C domain. Both residues are located in proximity of the spin label when the substrate hairpin is bound to the active site in the closed state (Fig. 7b). These PREs therefore suggest that the transiently formed, closed state interacts with the substrate hairpin in a similar manner as observed in the structure of the stably-closed hp-HP92/DbpA complex. In line with this finding, large PREs are observed for M161 and I355 upon stable formation of the closed state by addition of ADP/BeF_3_ (Fig. 7d, Figure 7 - figure supplement 1c). These PREs are also in good agreement with our structure of hp-HP92/DbpA complex and thereby provide an independent validation the this structure.

Overall, the PRE experiments demonstrate the transient formation of the closed state in the presence of HP92 and hp-HP92 RNAs, and also suggest that the substrate hairpin interacts with the transiently-formed active site in a similar manner as observed in the hp-HP92/DbpA structure. This hints at a pathway for the formation of the active state, where the closed state is transiently formed and, consequently, the substrate duplex is recruited to the pre-formed active site in a second step.

## Discussion

The crystallographic snapshots of the unwinding intermediates that we determined in this study fill a major gap in our understanding of the unwinding process of DEAD-box helicases and rationalize how the destabilization of the RNA duplex in the closed state is achieved. The structures can be readily integrated into a model of the unwinding process (Fig. 8): Prior to binding of the substrate duplex, DbpA mainly populates an open state, whereby the RecA_N domain tumbles independently from the other two domains. This open state, however, transiently adopts the closed-conformation. In the first step (1), the RNA duplex is recruited to this transiently-formed, closed-conformation. DbpA initially binds to 3 nt of the 5’ overhang and the first 3 nt of the substrate duplex, as depicted in the ds-HP92 complex structure (Fig. 4). Breathing of the duplex ends in the active site (step 2), which takes place on the μs-ms timescale (Snoussi and Leroy, 2001), leads to conformation 2 of the ss-HP92 complex (Fig. 5d). In this conformation, all residues still exhibit a continuous base stacking. Subsequently, in step (3), a transition to the canonical ssRNA product conformation (as observed in conformation 1 of the ss-HP92 complex) (Fig. 5c) prevents re-formation of the 3 base pairs. This strongly destabilizes the remaining duplex and increases the dissociation rate for the unbound RNA strand (step 4). The number of destabilized base pairs predicted based on these considerations is also in fair agreement with the number obtained from functional assays for the DEAD-box helicases eIF4A and Ded1 (2-3 and 4-5 base pairs, respectively (Raj et al., 2019; Rogers et al., 1999)).

**Fig. 8:**
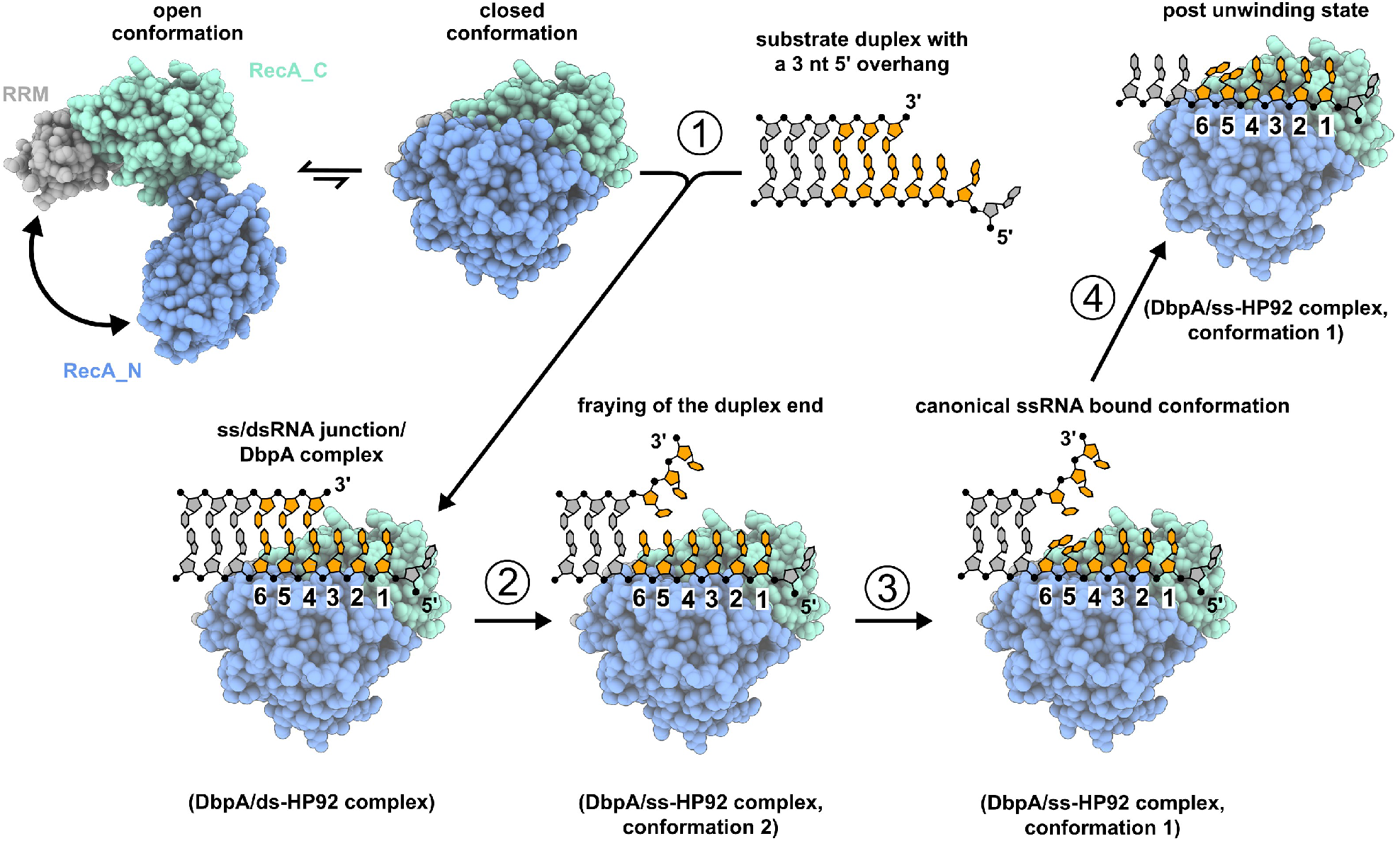
Model of the unwinding mechanism for duplexes with 5’ ssRNA overhangs. HP92 and ATP are omitted for clarity. The nucleotides that interact with the active site of DbpA are colored orange. DbpA transiently adopts the closed-conformation in the absence of substrate RNA (upper-left). (1) The closed-conformation binds to the ss/dsRNA junction, whereby nt 1-3 are single-stranded and nt 4-6 are still part of the duplex (as depicted in the structure of the DbpA/ds-HP92 complex). (2) Duplex breathing leads to a ssRNA-bound state with continuous base stacking (DbpA/ss-HP92 complex in conformation 2). (3) The nucleotides in positions 5/6 re-arrange to a conformation that is incompatible with duplex formation (DbpA/ss-HP92 complex in conformation 1). (4) The loss of these three base pairs leads to dissociation of the upper RNA strand.

Importantly, this model depicts only one of several different unwinding pathways that are compatible with our results. DbpA forms the closed state even more efficiently in the presence of ssRNA compared to the ss/dsRNA junction (Fig. 6c). It is therefore very likely that it can also bind more distal to the duplex such that only 1 or 2 nt of the duplex interact with the active site of the helicase core. Regarding blunt-end duplexes, fraying of the duplex ends will expose the 5’ ssRNA overhang that is necessary for binding. Moreover, in our experiments the attachment of the substrate duplex to HP92 enforces the interaction with the duplex end that is distant from the HP92 attachment site and prevents interaction with the proximal duplex end that carries a 3’ overhang. Our results therefore only provide insights into unwinding from the distant duplex end that carries the 5’ ssRNA overhang.

For most DEAD box helicases, the unwinding activity strongly increases with 3’ and with 5’ ssRNA overhangs relative to blunt end duplexes (Linder and Jankowsky, 2011), indicating that DEAD box helicases can also effectively initiate unwinding from 3’ ssRNA overhangs. This must proceed through a different mechanism, which might be analogous to the one described here and would involve binding to a ss/dsRNA junction with a 3’ ssRNA overhang. In this case, the duplex region would bind to positions 1-3, which are mainly formed by the RecA_C domain, and the 3’ ssRNA overhang would interact with positions 4-6. This notion is supported by the structure of the isolated RecA_C domain of the DEAD-box helicase Mss116 in complex with an RNA duplex, which shows that duplex binding to positions 1-3 is also possible (Mallam et al., 2012).

Interestingly, our ATPase assays with increasing 5’ overhangs indicate no specificity of DbpA for the ss/dsRNA junction over ssRNA, with 5’ ssRNA overhangs of increasing length being detrimental to the unwinding process. This is in agreement with previous reports on DbpA (Henn et al., 2010) and a similar observation has been made for the DEAD-box helicase eIF4A (Andreou et al., 2019). This finding supports a model where DEAD-box helicases act as ATP-fueled ssRNA binding proteins, rather than dsRNA-specific helicases. In this scenario, unwinding would be the result of stochastic binding to ss/dsRNA junctions, rather than a process that is targeted toward RNA duplexes. In line with this proposed lack of duplex-specificity, most DEAD-box helicases show strongly increased ATPase activity in the presence of ssRNA (Putnam and Jankowsky, 2013), indicating that interaction with a duplex is generally not necessary for the formation of the closed state. In addition, several DEAD-box helicases have been shown to act as ssRNA clamps (Ballut et al., 2005; Xiol et al., 2014) or in the remodeling of RNA/protein complexes (Bowers et al., 2006; Tran et al., 2007) and these two processes are unrelated to the interaction with dsRNA. In line with prior suggestions (Liu et al., 2008), we therefore favor a simple model, where the functions of DEAD-box helicases are explained by ATP-coupled, high-affinity ssRNA (or ss/dsRNA junction) binding.

In summary, our results for DbpA indicate that unwinding of duplexes with a 5’ ssRNA overhang proceeds through binding of the helicase core to the 5’ ss/dsRNA junction. The high conservation of the characteristic DEAD-box sequence motifs in DbpA (Figure 8 - figure supplement 1, (Putnam and Jankowsky, 2013)), together with the similarity of the ssRNA product-bound structure (ss-HP92 conformation 1) to the canonical product-bound state of DEAD-box helicases (Fig. 5c) suggests that our results can be generalized to other DEAD-box helicases. Nonetheless, in addition to the unwinding pathway via the 5’ ss/dsRNA junction described here, several other pathways are possible and remain to be described.

## Materials and Methods

### Protein expression and purification

For DbpA expression the gene coding for full length DbpA (UniProt accession code P21693) was PCR amplified from *E. coli* BL21(DE3) genomic DNA and cloned into a pETM-11 plasmid that codes for an N-terminal tobacco etch virus (TEV) protease cleavable hexahistidine tag. This plasmid was transformed into *E. coli* BL21(DE) codon plus cells. All growth media for protein expression were supplemented with 50 mg/L kanamycin and 34 mg/L chloramphenicol. Cells were grown in LB medium at 37 °C until an OD_600_ of 0.6-0.8 was reached. For production of unlabeled DbpA protein expression was induced at this stage by addition of 1 mM IPTG, the cells were shifted to 25 °C and harvested after 16-20 h. For production of ILMVA methyl group labeled DbpA 1 mL of the LB culture was used to inoculate 25 mL of H_2_O based M9 minimal medium and cells were grown to an OD_600_ of 0.6-0.8. Cells were harvested by centrifugation and used to inoculate 100 mL of D_2_O based M9-medium (containing 4 g/L deuterated glucose) at an OD_600_ of 0.15. Cells were grown over night at 37 °C and diluted 1/8 with fresh D_2_O based M9 medium (containing 2 g/L deuterated glucose) in the morning. Cells were grown at 37 °C until an OD_600_ of 0.8 was reached. The culture was shifted to 25 °C and 60 mg 2-Ketobutyric acid-(4-^13^C,3,3-d_2_), 100 mg 2-Keto-3- methyl-butyric acid-(dimethyl-^13^C_2_, 3-d) and 100 mg L-methionine-(methyl-^13^C) dissolved in 200 mL of D_2_O based M9 medium were added. After 45 min 100 mg L-alanine-(methyl-^13^C, 2-d) was added and 15 min later protein expression was induced by addition of 1 mM IPTG. Cells were harvested after 16-20 h and stored at -20 °C.

For purification the cell pellets were resuspended in 20 mL of Buffer A (400 mM NaCl, 50 mM sodium-phosphate, pH 7.4, 10 mM imidazole) supplemented with 0.1 % (v/v) triton-X-100, 1 mg/mL lysozyme and 0.1 mg/mL DNase I. Cells were lysed by sonication and cell debris was removed by centrifugation (30 min, 18000 g, 4 °C). The supernatant was applied to a gravity flow Ni-NTA column equilibrated in buffer A. The column was washed with 20 mL of buffer A, 10 mL of 1 M NaCl, 25 mM sodium-phosphate, pH 7.4 and 20 mL of buffer A + 10 mM imidazole. DbpA was eluted with buffer B (400 mM NaCl, 300 mM imidazole, 50 mM sodium-phosphate, pH 7.4). The N-terminal His-tag was removed by TEV protease digestion during dialysis against dialysis buffer (150 mM NaCl, 25 sodium-phosphate, pH 7.4, 1 mM DTT) over night at 4 °C. The TEV protease contained an N-terminal hexahistidine tag and was removed by a second Ni-NTA column equilibrated in dialysis buffer. The column was washed with dialysis buffer. Flow through and wash fractions were combined and ½ volume of 60 % (v/v) glycerol solution was added. DbpA was further purified using a 5 ml HiTrap HP Heparin column (GE healthcare) equilibrated in 100 mM NaCl, 20 mM HEPES, pH 7.3, 20 % (v/v) glycerol. DbpA was eluted using a linear NaCl gradient from 100 mM to 500 mM over 50 ml. DbpA containing fractions were pooled, concentrated and subjected to a size excluusion chromatography (SEC) step using a Superdex 75 16/600 column (GE healthcare) equilibrated in SEC buffer (125 mM NaCl, 25 mM HEPES, pH 7.3, 1 mM DTT). DbpA concentrations were determined based on the OD_280_ using an extinction coefficient of 26900 M^-1^cm^-1^.

### RNA production

RNAs were produced by in vitro transcription (IVTC) using homemade T7 polymerase (P266L mutant (Guillerez et al., 2005)). Single stranded DNA oligonucleotides, to which a DNA oligonucleotide corresponding to the T7 promotor sequence was hybridized, were used as templates. IVTCs were performed in 5-10 ml reactions containing 1 μM template DNA, 4 mM of each NTP, 15-50 mM MgCl_2_ (optimized for each RNA construct), 50 mM Tris, pH 8.0, 1 mM spermidine, 5 mM DTT, 0.01 % (v/v) triton X-100, 40 ug/ml T7 polymerase and were incubated at 37 °C for 3 h. Subsequently, 50 mM EDTA, pH 8.0 was added and the RNA was precipitated by addition of 0.7 vol of isopropanol. RNAs were purified using a DNAPac 100 column (22 x 250 mm, Dionex) heated to 70 °C. Buffers for purification contained 5 M Urea, 20 mM Tris, pH 8.0 (pH adjusted at RT) and 0 (buffer A) or 2 M NaCl (buffer B). RNAs were eluted using linear gradients from 5-15 % B to 20-40 % B depending on the RNA construct at a flowrate of 10 ml/min. Fractions containing the desired RNA product were pooled and precipitated by addition of 1 vol of isopropanol and incubation at -20 °C for at least 1 h. After centrifugation (8000 g, 4 °C, 45 min) the RNA pellet was washed with 75 % (v/v) ethanol, dried and resuspended in H _2_O. RNA concentrations were determined based on OD_260_. Extinction coefficients were calculated using the OligoAnalyzer Tool (IDT). See Appendix 1 – table 1 for the sequences of the RNA constructs used in this study.

### NMR experiments

All NMR experiments were conducted on Bruker 600 MHz and 800 MHz Avance Neo spectrometers equipped with nitrogen (600 MHz) or helium cooled (800 MHz) cryoprobes. Methyl-TROSY spectra were recorded using the SOFAST-HMQC pulse sequence (Schanda et al., 2005). Measurements were performed at 25 °C in SEC buffer supplemented with 5 % (v/v) D_2_O for the lock. DbpA concentrations in titration experiments with ILMVA-labeled DbpA were 40-70 μM and RNAs were added in 1.3 times excess over DbpA. For experiments where RNAs where titrated with DbpA, RNA concentrations were 100 μM and unlabeled DbpA was added in 1.3 times excess over RNA. To form the ADP/BeF_3_ bound state 2 mM MgCl_2_, 2 mM BeF_2_, 2 mM ADP and 10 mM NaF were added. CSPs were calculated according to the following equation

### CSP = ((ΔC/4)^2+(ΔH)^2))^0.5

(ΔC, ΔH chemical shift differences in ppm in the ^13^C and ^1^H dimensions).

Methyl group assignments in the free and RNA bound states of DbpA were published previously (Wurm, 2020; Wurm et al., 2021). Assignments for the RNA bound state could be partially transferred to the closed state (bound to ss-HP92 RNA and ADP/BeF_3_) based on 3D-NOESY spectra in combination with the structure of the DbpA/ss-HP92/ADP/BeF_3_ complex. The sample for the NOESY spectra contained 250 μM ILMVA methyl group labeled DbpA, 350 μM ss/HP92 RNA, 110 mM NaCl, 25 mM Arg/Glu, 25 mM HEPES, pH 7.3, 1 mM DTT, 5 mM MgCl_2_, 2 mM ADP, 4 mM BeF2 and 15 mM NaF. SOFAST-HMQC based 3D-CCH- and 3D-HCH-NOESY spectra (Rossi et al., 2016) were recorded with a mixing time of 250 ms.

Imino proton signals of the substrate duplex of the hp-HP92 RNA in the free state were assigned using a 2D ^1^H-^1^H NOESY spectrum (mixing time 130 ms) recorded in SEC buffer at 10 °C at an RNA concentration of 200 μM. The assignment of the imino protons of HP92 was published previously (Wurm et al., 2021).

NMR spectra were processed with Topspin 4.0.2 or NMRPipe 9.6 (Delaglio et al., 1995) and analyzed using NMRPipe and CARA (Keller, 2004).

### PRE measurements

RNAs for site selective spin labelling contained a 4-thiouridine at the desired labelling position and were chemically synthesized by Dharmacon, see Appendix 1 – table 1 for RNA sequences. RNAs were deprotected according to the protocol provided by Dharmacon. For spin labelling 100 μM of the respective RNA was incubated at room temperature in the dark for 24 h with a 100-fold excess of 4-(2-Iodoacetamido)-TEMPO in a buffer containing 20 % (v/v) DMSO and 100 mM HEPES, pH 8.0. Unreacted 4-(2-Iodoacetamido)-TEMPO was removed by two sodium acetate/EtOH precipitation steps. PRE experiments were performed in NMR buffer lacking DTT using a 1.3 times excess of RNA over DbpA. DbpA concentrations ranged from 40 to 60 μM. SOFAST-HMQC spectra were recorded before and after reduction of the spin label by addition of 2 mM sodium ascorbate. Peak volumes in both spectra were integrated with NMRPipe and the PRE was calculated as the ratio of the peak volumes before reduction (I) divided by the peak volumes after reduction (I_0_). PRE values for the geminal methyl groups of Leu and Val residues were averaged.

### Helicase assays

Fluorescence based single turnover helicase assays were conducted at 25 °C in 96-well plates using a TECAN spark platereader. The assay takes advantage of the fluorescence quenching effect of the 5’ guanosine residues of the 9mer RNA on the 5’ fluorescein label. This effect is strongly reduced, when the labeled 9mer is hybridized to the HP92 containing RNA and leads to a reduction of the fluorescence intensity upon unwinding. Rebinding of the 9mer RNA to HP92 containing RNA is prevented by an excess of unlabeled 9mer RNA. See Appendix 1 – table 1 for RNA sequences. 5’ fluorescein labeled RNAs were obtained from IDT. Reaction mixtures (100 uL) contained 125 mM NaCl, 25 mM HEPES, pH 7.3, 2 μM DbpA, 5 mM MgCl_2_, 25 nM 5’ fluorescein labeled 9mer RNA, 37.5 nM HP92 containing RNA, 2 μM unlabelled 9mer RNA, 3 mM of the respective nucleotide (ADP, ATP, ADPNP, ADPCP or ATPγS). For the ADP/BeF_3_ condition 3 mM BeF_2_ and 10 mM NaF were added in addition to ADP. Initially, 5’ fluorescein labeled 9mer RNA (at 1 μM) and HP92 containing RNA (at 1.5 μM) were hybridized in SEC buffer supplemented with 5 mM MgCl_2_ by heating to 95 °C and subsequent cooling to room temperature over ∼ 1 min. Then all components except for the nucleotide were mixed and preincubated for 5 min in the platereader. The unwinding reaction was started by addition of nucleotide (+ BeF_2_ and NaF in the case of the ADP/BeF_3_ condition) and the fluorescein fluorescence (excitation/emission filter wavelength 485 nm/535 nm) was recorded every 30 s. Fluorescence time courses were fitted to the following equation

F = F_0_ + ΔF·e^-k·t^

(F fluorescence intensity, F_0_ basal fluorescence intensity, ΔF fluorescence intensity difference due to unwinding, k unwinding rate, t time) using Matlab. All helicase assays were performed in triplicate using the same DbpA preparation, which was stored in small aliquots at -80 C until use. The reported values are mean and standard deviation of the three measurements.

### ATPase assays

The ATPase activity of DbpA in the presence of different RNA constructs was determined using a coupled pyruvate kinase/lactate dehydrogenase assay (Kiianitsa et al., 2003; Tsu and Uhlenbeck, 1998). The production of ADP is coupled to NADH oxidation, which can be monitored based on NADH absorption at 340 nm. Reaction mixtures (150 uL) contained 0.2-1 μM DbpA, 3 mM ATP, 5 μM HP92 containing RNA, 0 or 6.5 μM 9mer RNA, 450 μM NADH, 1.5 mM pyruvate, 10 u/ml pyruvate kinase/lactate dehydrogenase (from rabbit muscle, Sigma Aldrich, #P0294) and 5 mM MgCl_2_ in SEC buffer. Absorption measurements at 340 nm were conducted in 96-well plates every 20 s using a TECAN spark platereader at 25 °C. ATPase rates were determined based on the following equation after linear fitting of the linear region of the absorption time courses k_obs_= - dA/dt*(1/K_path_)*(1/[DbpA]) (k_obs_ ATPase activity, dA/dt slope of the linear fit to the absorption time course, K_path_ molar absorption coefficient of NADH at 340 nm for the path length in the 96-well plate (2.22 mM^-1^), [DbpA] DbpA concentration). All ATPase assays were performed in triplicate using the same DbpA preparation, which was stored in small aliquots at -80 C until use. The reported values are mean and standard deviation of the three measurements.

### Structure modeling

Structure modeling was performed in torsion angle space using CYANA 3.98 (Güntert et al., 1997). The 1:1 complex between hp-HP92 RNA and DbpA was modeled based on chain B (DbpA), nt 1- 22 of chain F and nt 26-42 of chain G. The torsion angles in these residues were fixed (except for the ε backbone angle of nt 22 and the α, β, δ and ζ backbone angles of nt 26) and nt 23-25 of the hp- HP92 were modeled to connect chains F and G using the regularize macro implemented in CYANA. 50 structures were calculated and the structure with the lowest target function was chosen to represent the model.

To model the elongated duplex of the ds-HP92 RNA (Fig. 4) two cytosine residues (corresponding to nt 45 and 46) were added to the 5’ end of the ds-HP92 RNA. For these cytosines A-form helix restraints (Richardson et al., 2008) and Watson-Crick hydrogen bond restraints to nt G2/G3 were introduced. All angles of the ds-HP92 RNA except for the δ and ε backbone angles of nt 44 were fixed. 50 structures were calculated by simulated annealing and the structure with the lowest target function was chosen to represent the model.

### Analytical size exclusion chromatography

Analytical size exclusion chromatography of RNA and RNA/DbpA complexes was performed at 25 °C using an XBridge Protein BEH SEC 200Å Column (7.8 mm x 300 mm) and SEC buffer (flowrate 0.5 ml/min). RNAs were folded by heating the RNAs at a concentration of 80 μM in SEC buffer to 95 °C for 1 min, followed by rapid cooling on ice. 5 uL of solution containing RNA at a concentration of 40 μM in SEC buffer were injected for each run. DbpA was added at a 1.3 fold excess and 2 mM ADP, 2 mM BeF_2_, 2 mM MgCl_2_ and 10 mM NaF were added to form the RNA/DbpA/ADP/BeF_3_ complexes. The RNA/DbpA/ADP/BeF_3_ complex was incubated at least 20 min at room temperature before analysis. RNA elution was monitored by recording the absorption at 260 nm.

### Crystallization and structure determination

The DbpA/RNA/ADP/BeF3 complexes for crystallization were prepared in SEC buffer using DbpA and RNA concentrations of 200 and 260 μM, respectively. Before mixing with DbpA RNAs were folded at a concentration of 700 μM in H_2_O by heating to 95 °C for 1 min followed by rapid cooling on ice and subsequent addition of NaCl to a final concentration of 125 mM. Then 3 mM ADP, 5 mM BeF_2_, 5 mM MgCl_2_ and 16 mM NaF were added and the complex was incubated at least 1 h at room temperature before crystallization. Crystals were grown using the hanging drop vapor diffusion method at 20 °C after mixing 1 ul of the complex solution with 1 ul of the reservoir solution. Crystals appeared after 1 to 2 days. Crystals were cryoprotected using reservoir solution + 20 % (v/v) PEG400 and subsequently flash frozen in liquid nitrogen. The following reservoir solutions were used:

hp-HP92 RNA: 200 mM NH_4_/tartrate, 100 mM Tris, pH 8.0, 20% (w/v) PEG3350

ds-H92 RNA: 0.2 M di-Sodium tartate, 20 % (w/v) PEG 3350

ss-HP92 RNA: 400 mM KSCN, 100 mM Tris, pH 8.0, 20 % (w/v) PEG 3350

Diffraction data were collected at −170 °C and at a wavelength of 0.9766 Å at the EMBL/DESY beamline P13 (Hamburg, Germany) (Cianci et al., 2017) (for the ss/HP92 RNA/DbpA complex) or at a wavelength of 1.000 Å at the SLS beamline X06SA (Villigen, Switzerland) (Mueller et al., 2012) (for the hp/HP92 and ds/HP92 RNA/DbpA complexes). The collected data was integrated, merged and scaled using XDS (Kabsch, 2010). Phasing of the hp-HP92 RNA complex was performed by molecular replacement using PHASER (McCoy et al., 2007) and the structures of the DEAD-box helicase VASA (PDB 2DB3) (Sengoku et al., 2006) and of the RRM of the DbpA homolog YxiN (PDB 3MOJ) (Hardin et al., 2010) as search models. For the other two complexes the DbpA structure of the hp-HP92 RNA complex was used as a molecular replacement search model. The structures were refined by iterative rounds of manual model building in COOT (Emsley and Cowtan, 2004) and refinement using Phenix.refine (Afonine et al., 2012). Data collection and refinement statistics are summarized in Figure 3 - figure supplement 1. Structure representations were generated with UCSF ChimeraX (Pettersen et al., 2021).

## Data availability

Structure factors and atomic coordinates have been deposited in the PDB with accession codes 7PLI (ss-HP92/DbpA complex), 7PMQ (hp-HP92/DbpA complex) and 7PMM (ds-H92/DbpA complex).

## Acknowledgment

I am indebted to Remco Sprangers for his generous support, fruitful discussion, lab space and spectrometer time. I would like to thank Jens Wöhnert and Konstantin Neißner for acquisition of the diffraction data of the ss-HP92 RNA/DbpA complex and Olga Rudi for help with protein preparation. The Paul Scherrer Institut, Villigen, Switzerland and the EMBL Hamburg are acknowledged for synchrotron radiation beamtime at beamline P13 at the PETRA III storage ring and at beamline PXI of the SLS. I am grateful to Christoph Engel and Ursula Neu for helpful discussions regarding structure refinement and I would like to thank Johanna Stöfl for lab support and Jan Overbeck and Jobst Liebau for critical reading of the manuscript. The Deutsche Forschungs Gemeinschaft is acknowledged for funding (DFG grant no. WU 988/1-1).

## Appendix

**Figure 1 - figure supplement 1:**
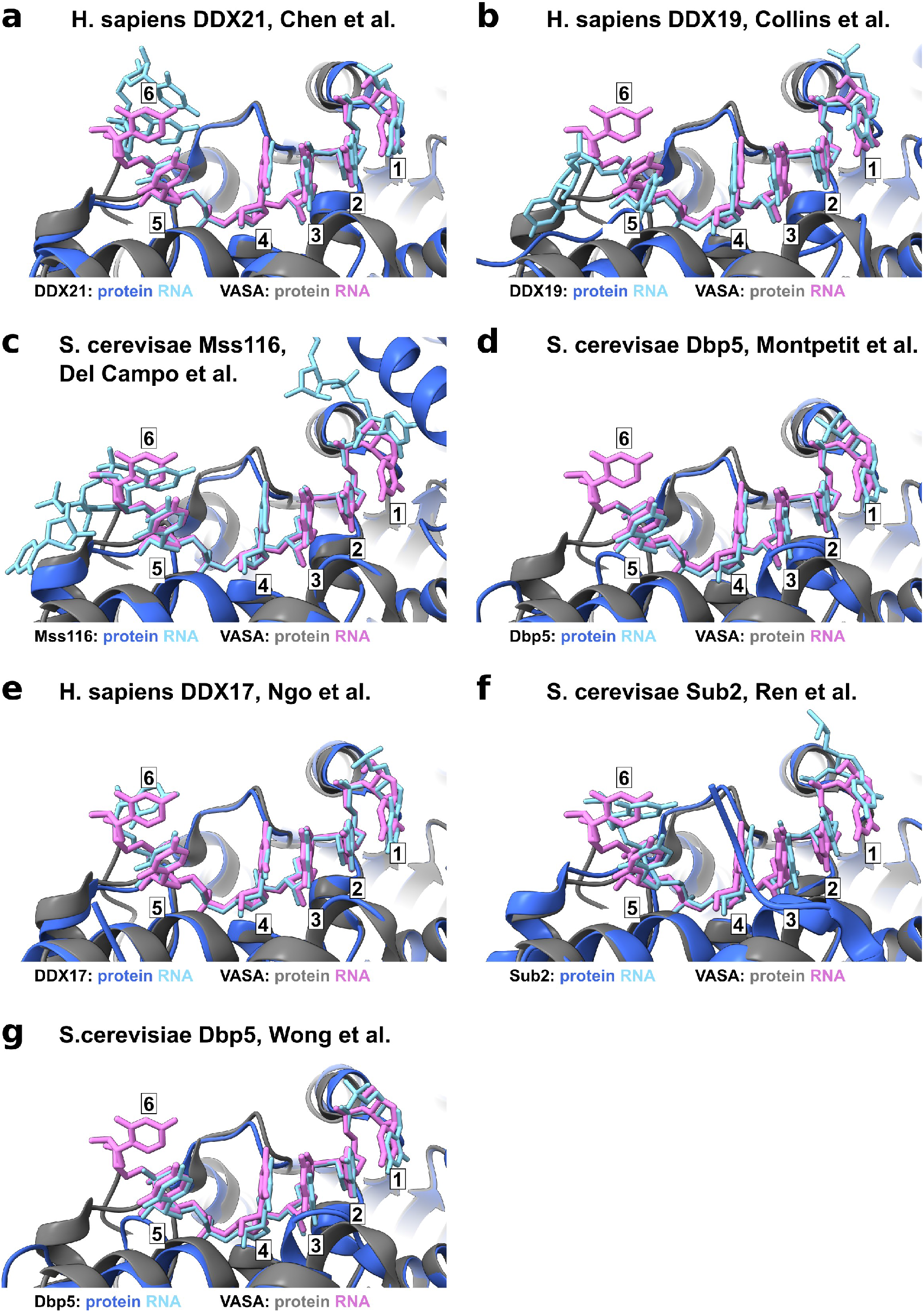
Published structures of ssRNA product-bound DEAD-box helicases in the closed-conformation reveal a highly conserved binding mode. Close-up of the active site of the helicase core bound to ssRNA. Shown are comparisons between the structure of the DEAD-box helicase VASA from *D. melanogaster* with the structures of *(a) H. sapiens* DDX21 *(Chen et al., 2020),* (b) *H. sapiens* DDX19 (Collins et al., 2009), (c) *S. cerevisiae* Mss116 (Del Campo and Lambowitz, 2009), (d) *S. cerevisiae* Dbp5 (Montpetit et al., 2011), (e) *H. sapiens* DDX17 (Ngo et al., 2019), (f) *S. cerevisiae* Sub2 (Ren et al., 2017) and *S. cerevisiae* Dbp5 (Wong et al., 2016). The VASA helicase and the bound ssRNA are shown in grey and pink. The other helicases and bound ssRNAs are colored dark and light blue. The nucleotides that interact with the active site are numbered 1-6. Basically identical conformations are observed for nt 1-5 and the phosphate group of nt 6, whereas the conformation of the ribose and base of nt 6 are less well conserved.

**Figure 2 - figure supplement 1:**
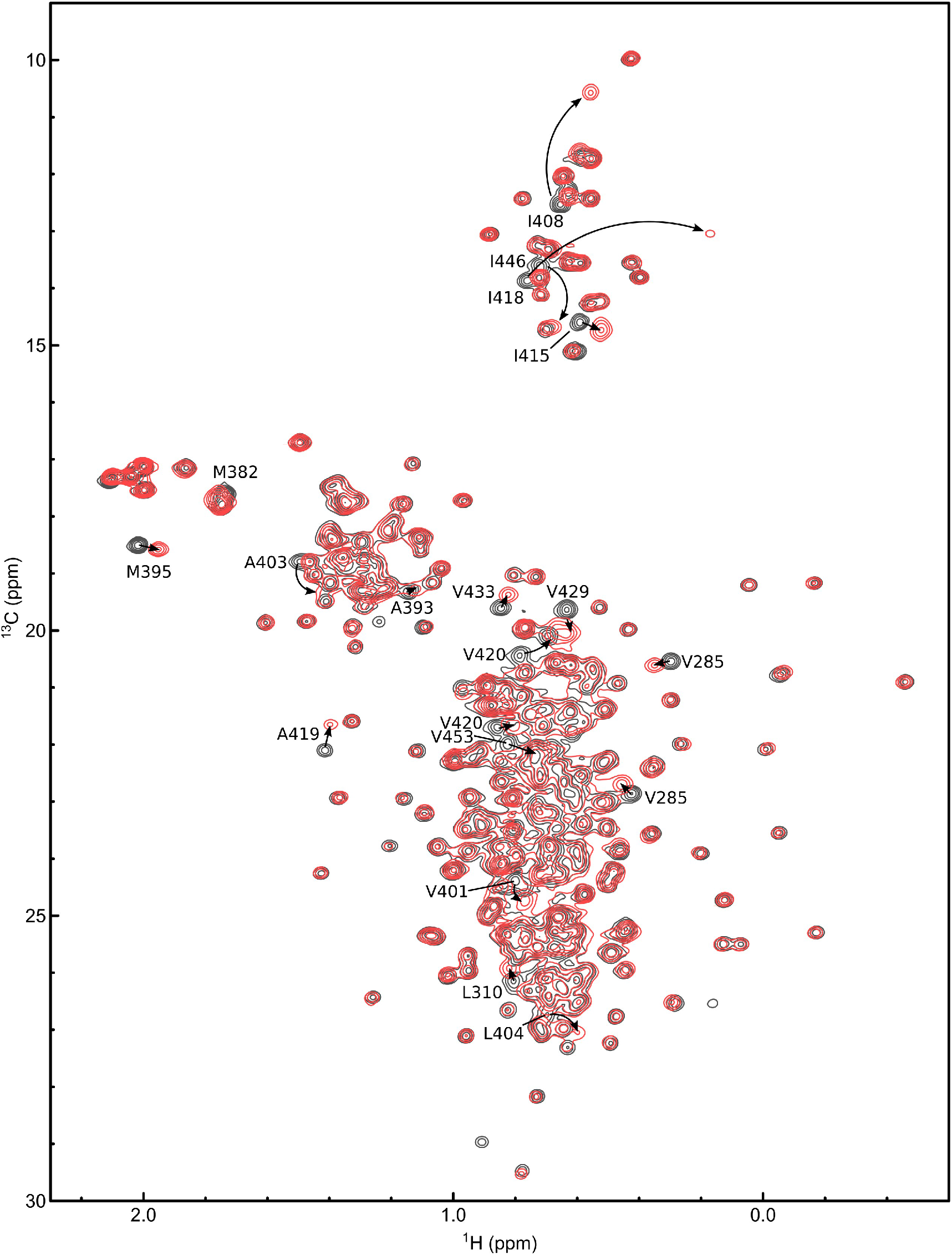
ATPγS is rapidly hydrolyzed by DbpA in the presence of ss- HP92 RNA. (**a**) Overlay of ^1^H^13^C-HMQC spectra of ILMVA-methyl group labeled DbpA in complex with ss- HP92 RNA after addition of ADP (red) or ATPγS (blue). The spectra are virtually identical indicating that the ATPγS has been hydrolyzed to ADP before the start of the measurement. (**b**) Overlay of ^1^H^13^C-HMQC spectra of ILMVA-methyl group labeled DbpA after addition of ADP (red) or ATPγS (blue). In the absence of ss-HP92 the spectra are clearly different indicating that ATPγS is only hydrolyzed in the presence of ss-HP92 and that binding of ADP and ATPγS can be discriminated based on ^1^H^13^C-HMQC spectra of DbpA.

**Figure 2 - figure supplement 2:**
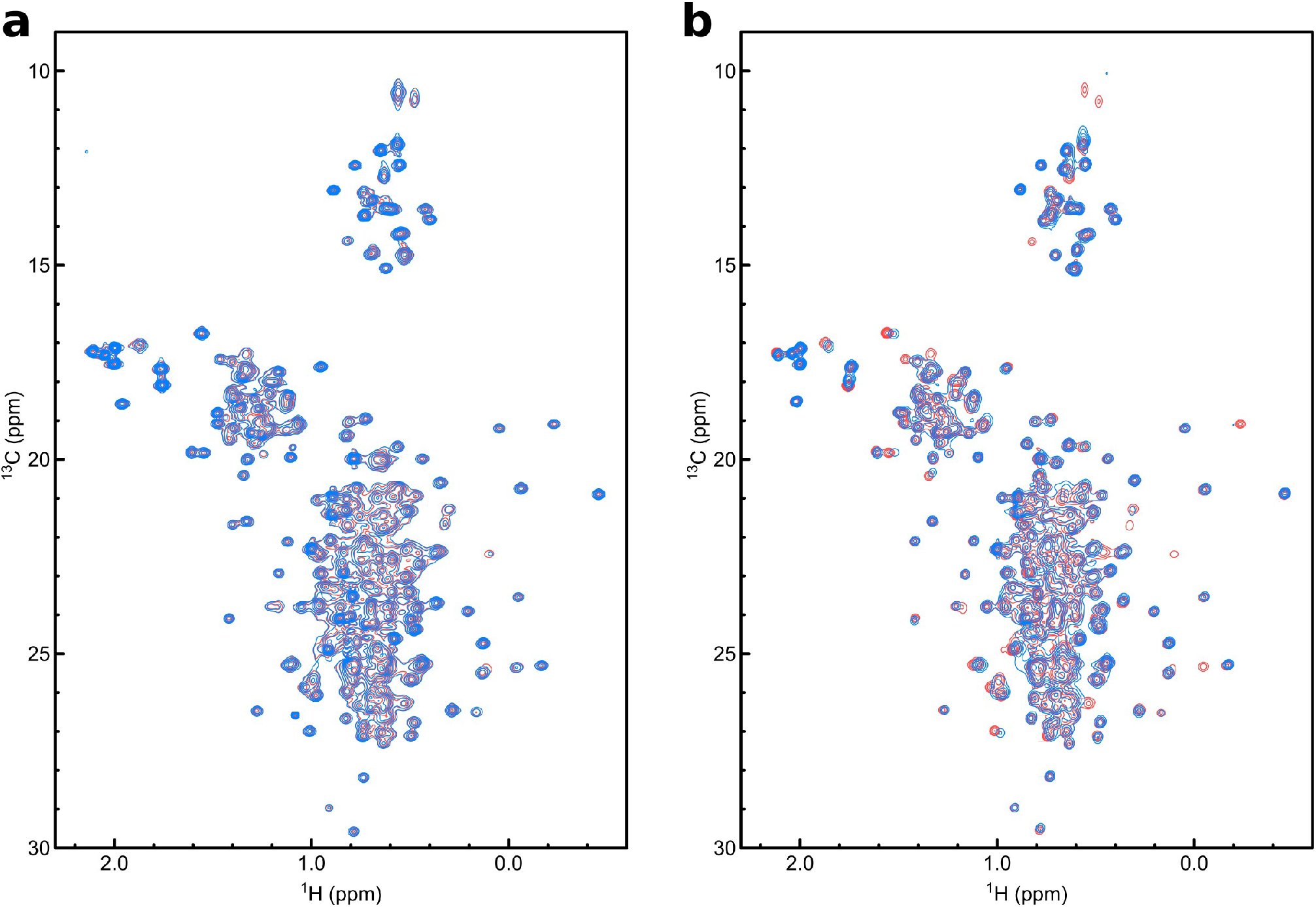
Binding of hp-HP92 to DbpA followed by methyl NMR. Overlay of ^1^H^13^C-HMQC spectra of ILMVA-methyl group labeled DbpA in the apo state (black) and after addition of a 1.3-fold excess of hp-HP92 RNA (red). Assignments are given for methyl group signals that show large CSPs and the peak position in the complex is indicated by an arrow. Note that all signals that show large CSPs belong to the RRM (res. 380-457) or are located in the proximity of the RRM (V285/L310). See figure 2 of the main text for quantification of the CSPs.

**Figure 2 - figure supplement 3:**
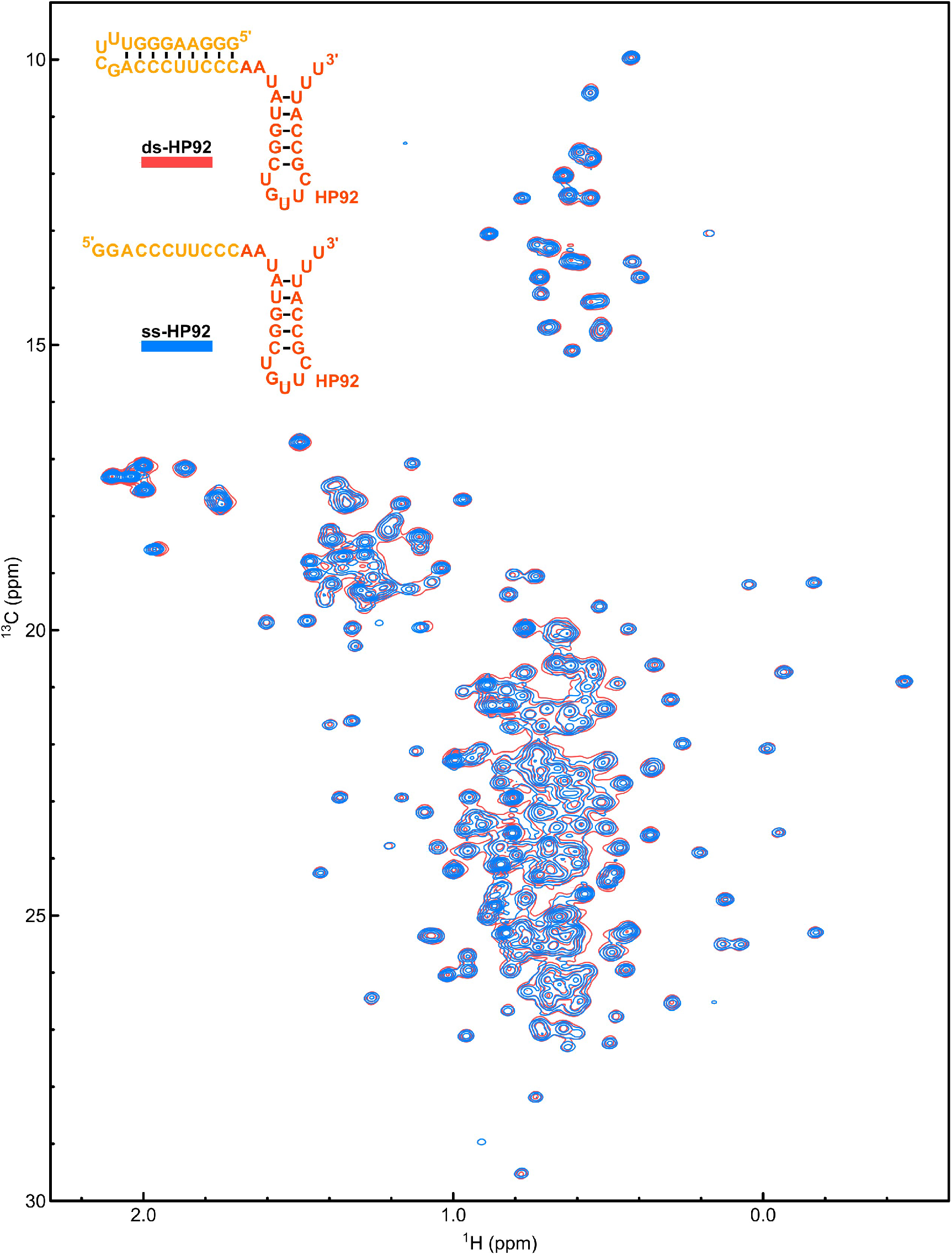
Spectra of DbpA in complex with ss-HP92 and hp-HP92 RNAs are virtually identical. Overlay of ^1^H^13^C-HMQC spectra of ILMVA-methyl group labeled DbpA in complex with the hp- HP92 (red) or the ss-HP92 RNA (blue). The spectra are virtually identical (see also Fig. 2 of the main text). The RNA constructs are shown in the top left.

**Figure 2 - figure supplement 4:**
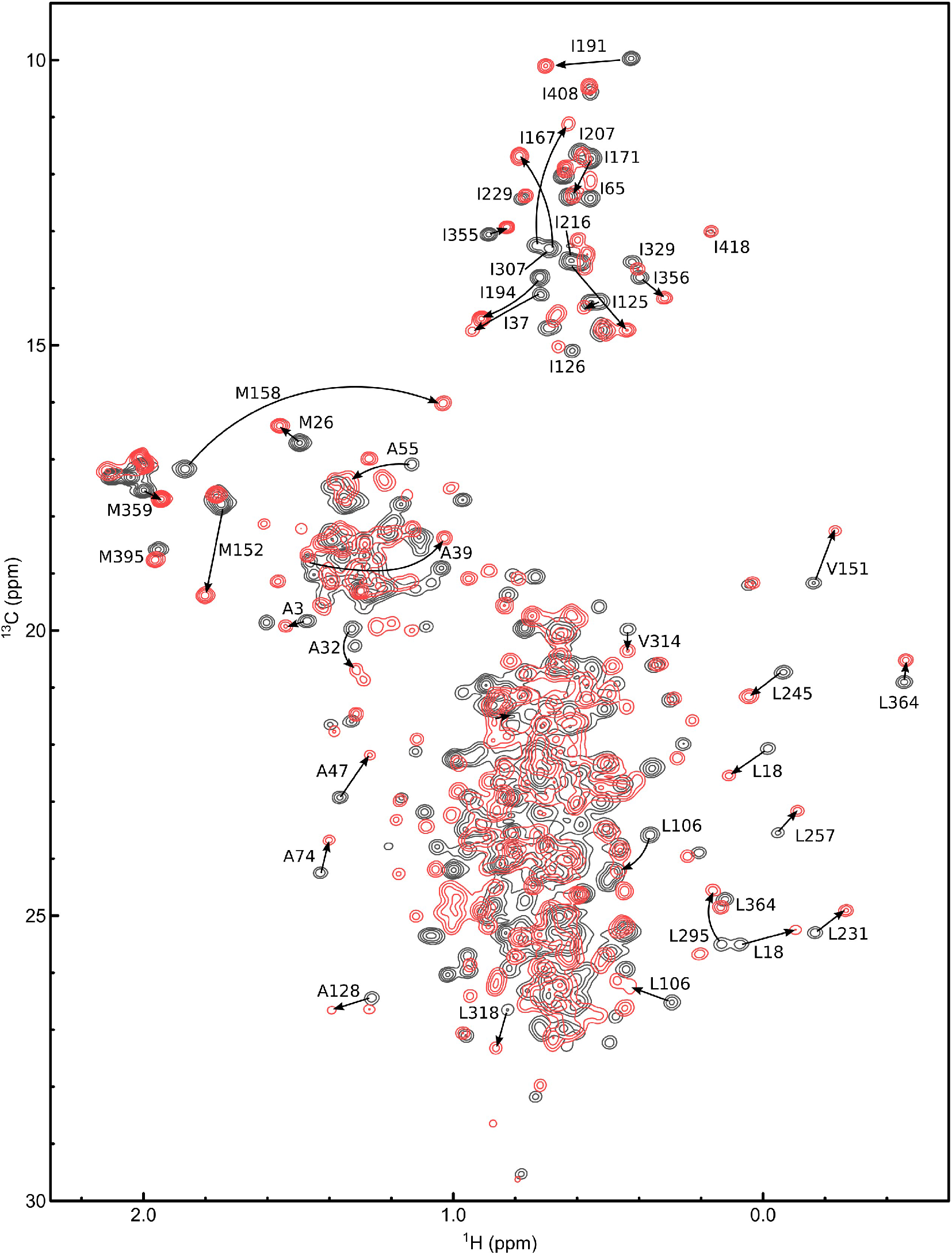
Binding of ADP/BeF_3_ to the hp-HP92/DbpA complex followed by methyl NMR. Overlay of ^1^H^13^C-HMQC spectra of ILMVA-methyl group labeled DbpA in complex with hp-HP92 RNA (black) and after addition of a ADP/BeF_3_ (red). Assignments are given for methyl group signals that show large CSPs and the new peak position is indicated by an arrow. Note that signals from both RecA domains show large CSPs. See Fig. 2 of the main text for quantification of the CSPs. As shown previously (Wurm et al., 2021) these CSPs signify the formation of the closed state of the helicase core.

**Figure 2 - figure supplement 5:**
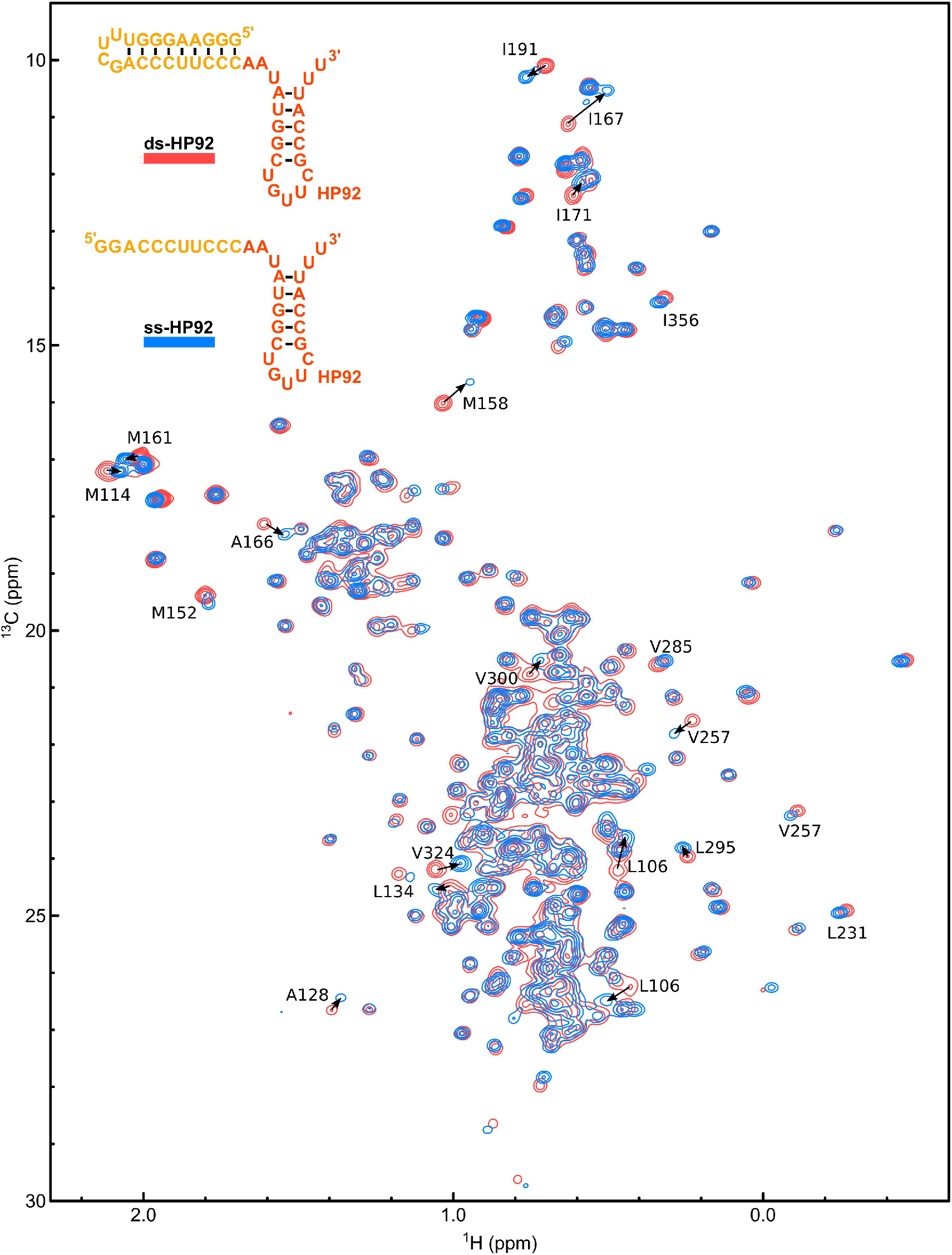
Spectra of DbpA/ADP/BeF_3_ in complex with and ss-HP92 and hp-HP92 complex are similar, but display marked differences. Overlay of ^1^H^13^C-HMQC spectra of ILMVA-methyl group labeled DbpA in complex with ADP/BeF_3_ and the hp-HP92 RNA (red) or in complex with ADP/BeF_3_ and the ss-HP92 RNA (blue). Assignments are shown for signals with pronounced differences between the two constructs. The differences cluster in the vicinity of the RNA binding site of the RecA domains. See Fig. 2 of the main text for a quantitative comparison. The RNA constructs are shown in the top left.

**Figure 3 - figure supplement 1:**
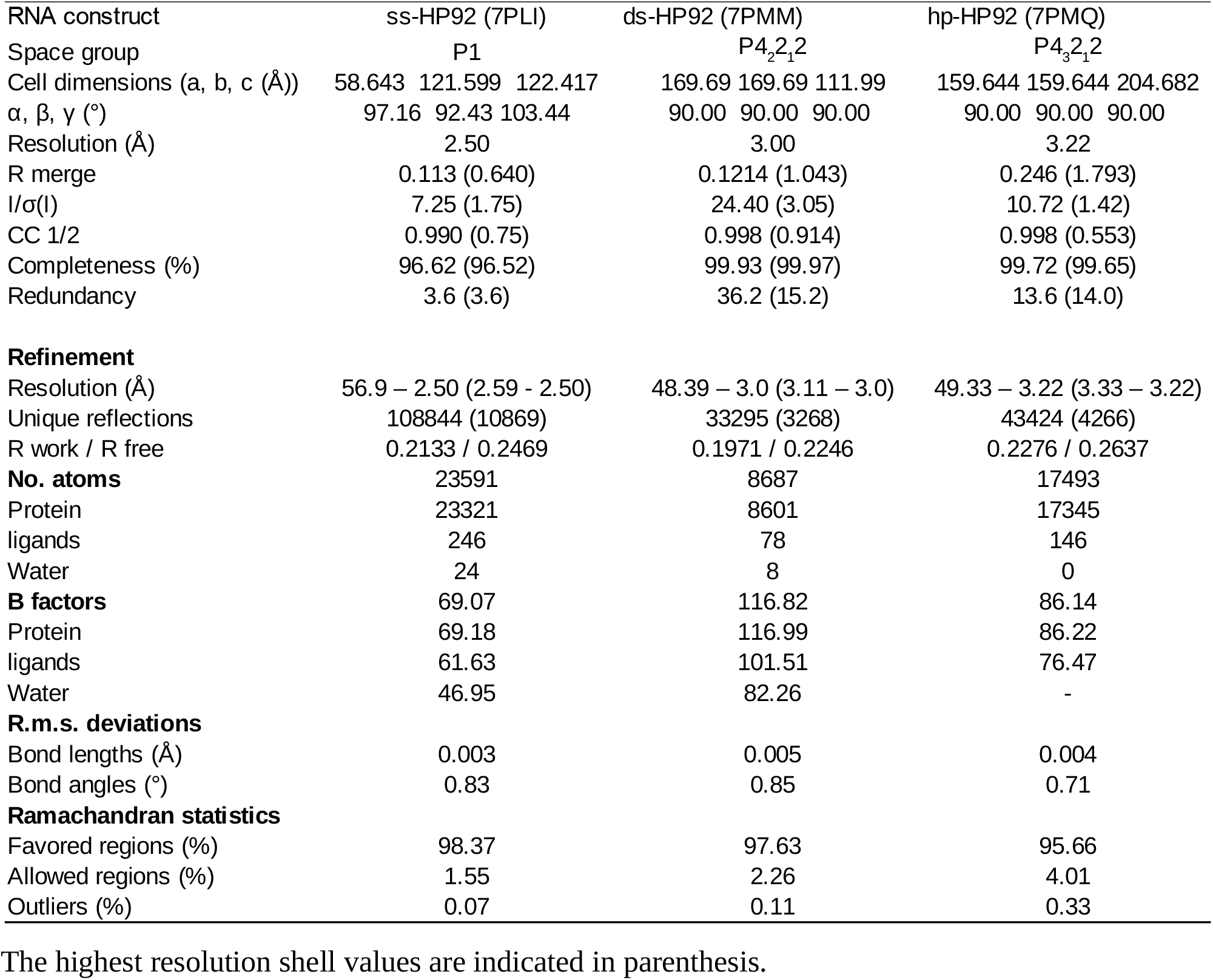
**Crystallographic data and refinement**

**Figure 3 - figure supplement 2:**
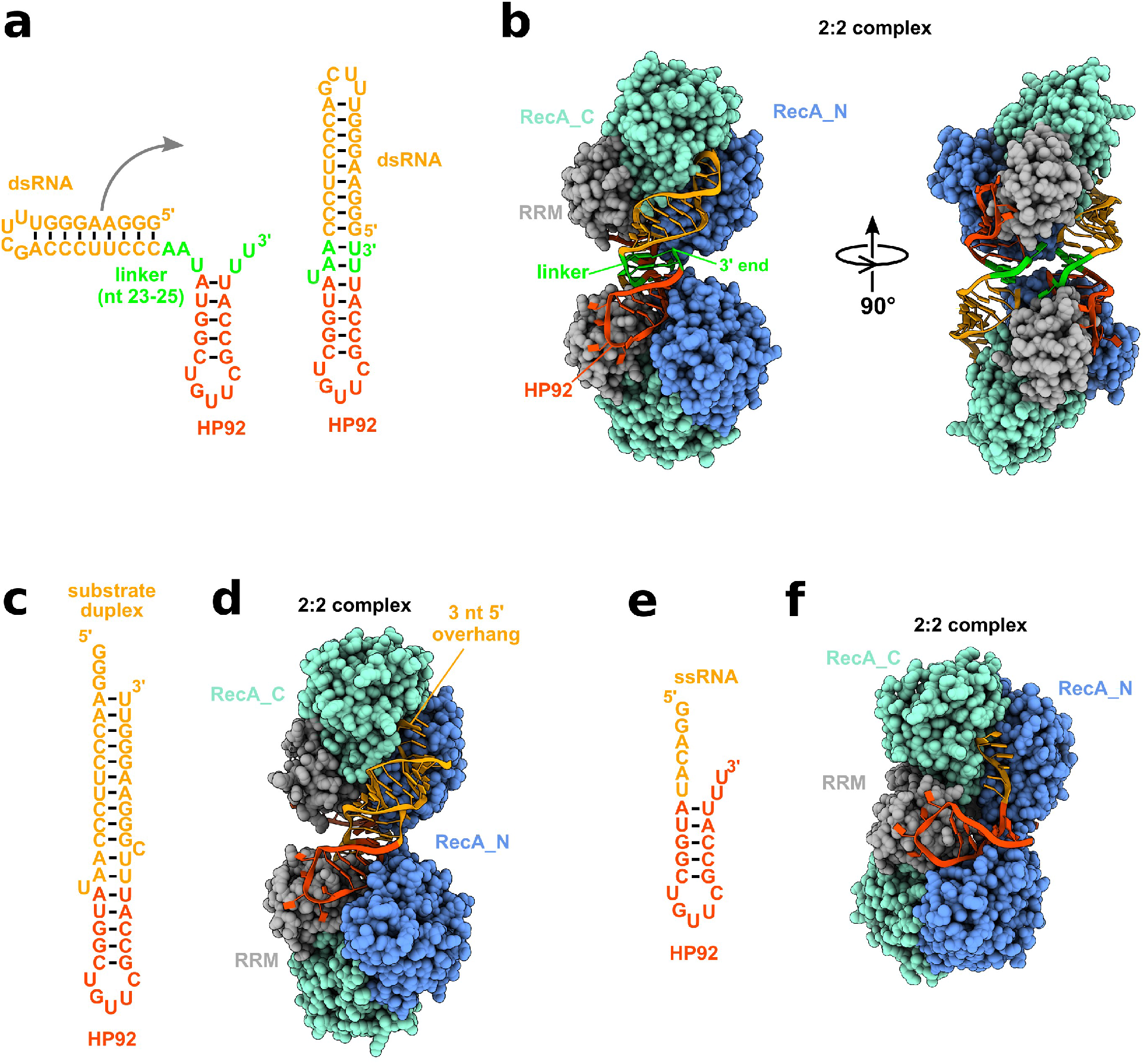
Crystallized RNA constructs and 2:2 complexes. (**a**) RNA construct used for crystallization of the hp-HP92/DbpA/ADP/BeF_3_ complex. The expected secondary structure of the hp-HP92 RNA is shown on the left and the secondary structure observed in the crystal is shown on the right. The linker (nt 23-25 green) between the substrate hairpin (orange) and HP92 (red) forms two base pairs with the uridines at the 3’ end, which leads to co coaxial stacking of the two helices. (**b**) Crystallized 2:2 complex between 2 DbpA molecules and 2 hp-HP92 RNAs (chains A/B/FG). Each RNA interacts with two DpbA molecules: HP92 binds to the RRM of one DbpA molecule and the substrate hairpin interacts with the RecA domains of the other DbpA molecule. (**c**) ds-HP92 RNA construct used for crystallization of the ds-HP92/DbpA/ADP/BeF_3_ complex. To facilitate crystallization the construct was designed to mimic the conformation of the hp-HP92 RNA observed in the crystal (compare to (a)). Instead of the hairpin the ds-HP92 RNA possess a ss/dsRNA junction with 5’ ssRNA overhang of 3 nt that interacts with the RecA domains of the helicase core. (**d**) Crystallized 2:2 complex between 2 DbpA molecules and two ds-HP92 RNAs (chains A/B/C/D). (**e**) ss-HP92 RNA construct used for crystallization of the ss-HP92/DbpA/ADP/BeF_3_ complex. (**f**) Crystallized 2:2 complex between two DbpA molecules and two ss-HP92 RNAs (chains A/B/C/D)

**Figure 3 - figure supplement 3:**
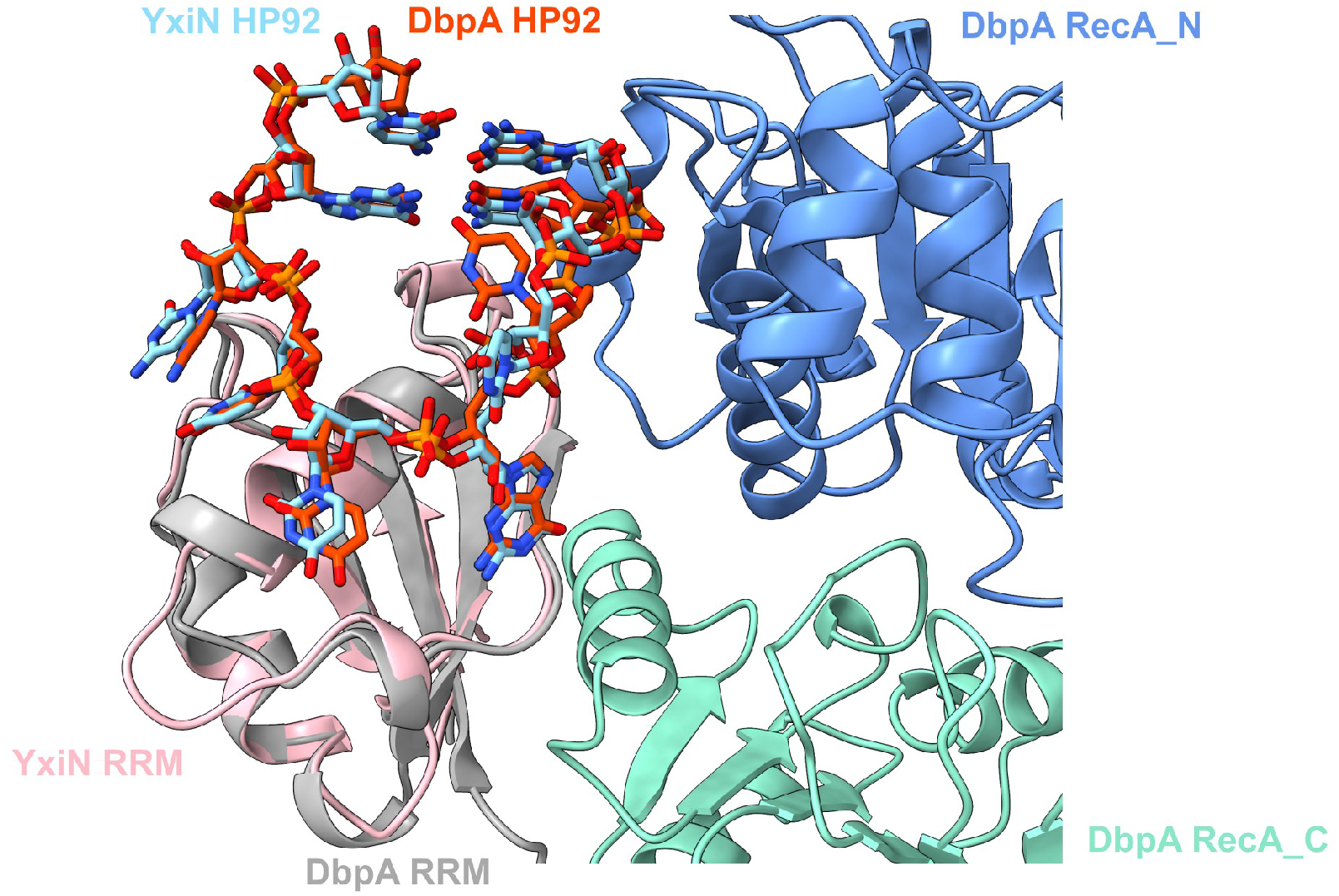
The HP92 RNA binding modes of the RRMs of DbpA and its *Bacillus subtilis* homolog YxiN are very similar. Overlay of the structures of the YxiN RRM bound to a fragment of the 23S rRNA (PDB ID 3moj) and the structure of the Hhp-HP92/DbpA/ADP/BeF_3_ complex (chains B/G). The RecA_figs3c. The bound HP92 RNAs are shown in stick representation (DbpA red, YxiN light blue). For clarity only nt 2550-2559 of the rRNA fragment and the corresponding nt of the hp-HP92 RNA are shown.

**Figure 3 - figure supplement 4:**
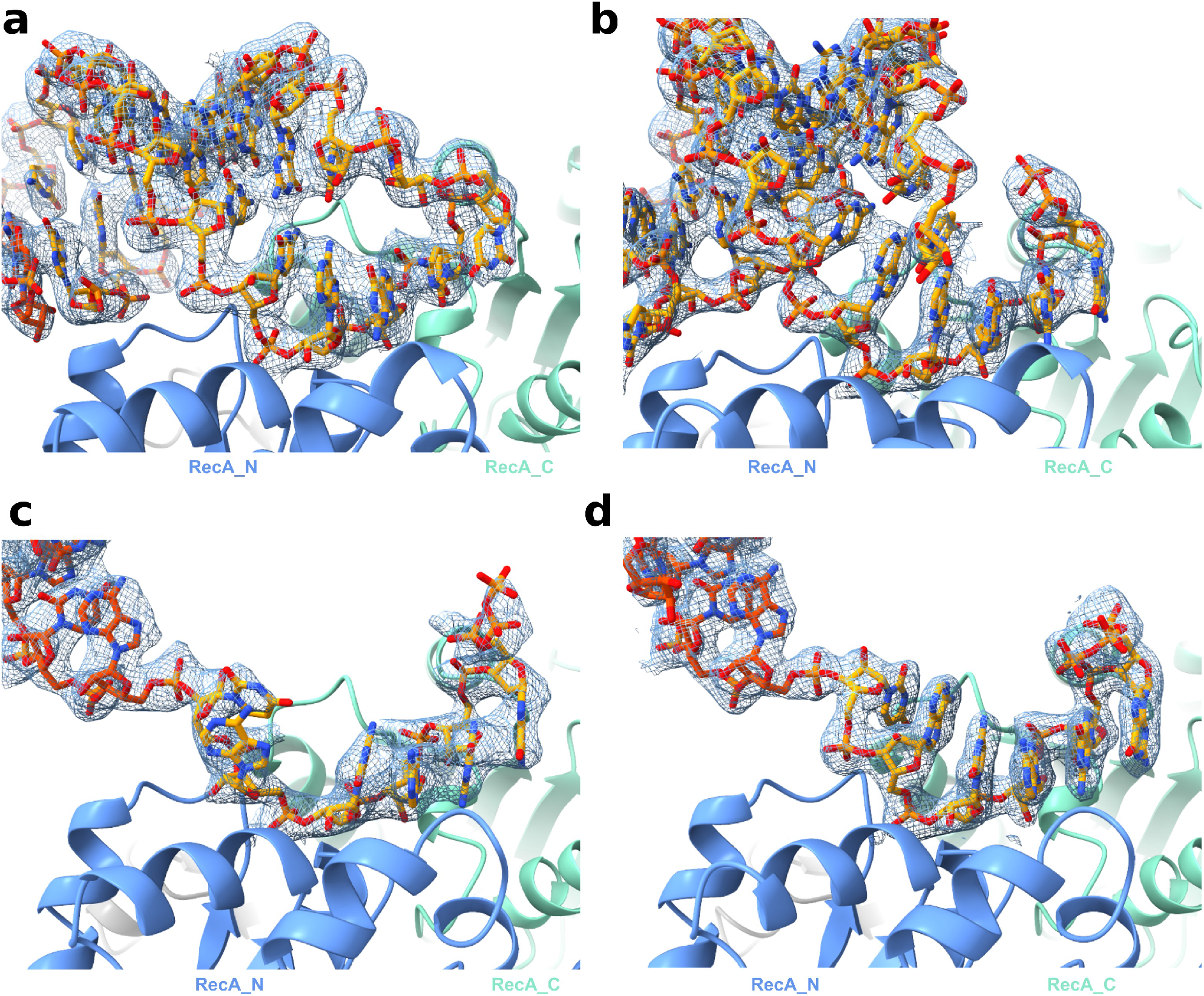
Electron density maps. Composite omit 2Fo - Fc maps contoured at 1.0 σ are shown as a blue mesh for the RNAs depicted in Fig. 3, 4 and 5 of the main text. (**a**) hp-HP92/DbpA/ADP/BeF_3_ complex (chain F). (**b**) ds- HP92/DbpA/ADP/BeF_3_ complex (chain D). (**c**) ss-HP92/DbpA/ADP/BeF_3_ complex conformation 1 (chain L). (**d**) ss-HP92/DbpA/ADP/BeF_3_ complex conformation 2 (chain C).

**Figure 5 - figure supplement 1:**
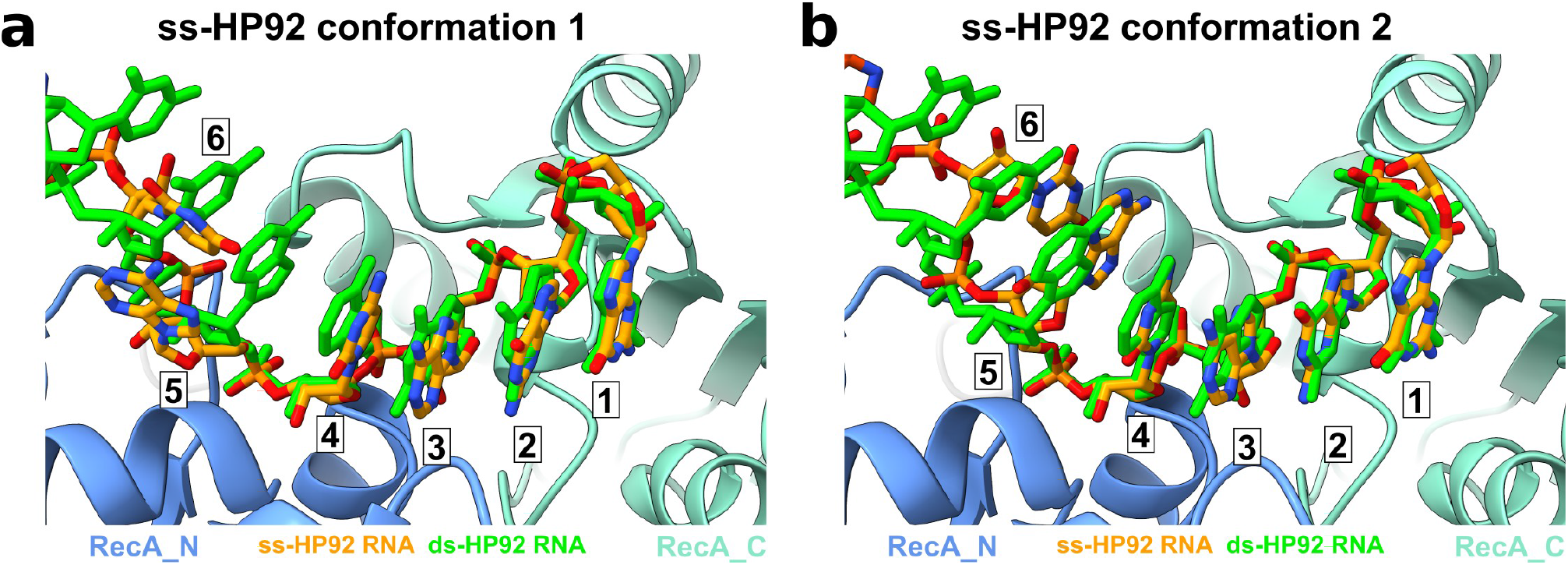
The ss-HP92 RNA conformation 2 is more similar to the ds- HP92 RNA conformation than ss-HP92 RNA conformation 1 Close-ups of the ss-HP92 RNA in (a) conformation 1 (chain L, orange) and (b) conformation 2 (chain C, orange) bound to the active site of DbpA. For comparison the ds-HP92 RNA is shown in green. For clarity the opposite strand that base pairs with nt 4-6 is omitted for ds-HP92 RNA. The nucleotides that interact with the active site are numbered 1-6. The conformation and base orientation of nucleotides 5 and 6 in conformation 2 is more similar to the ds-HP92 RNA conformation compared to conformation 1, where the bases are flipped by 90°.

**Figure 5 - figure supplement 2:**
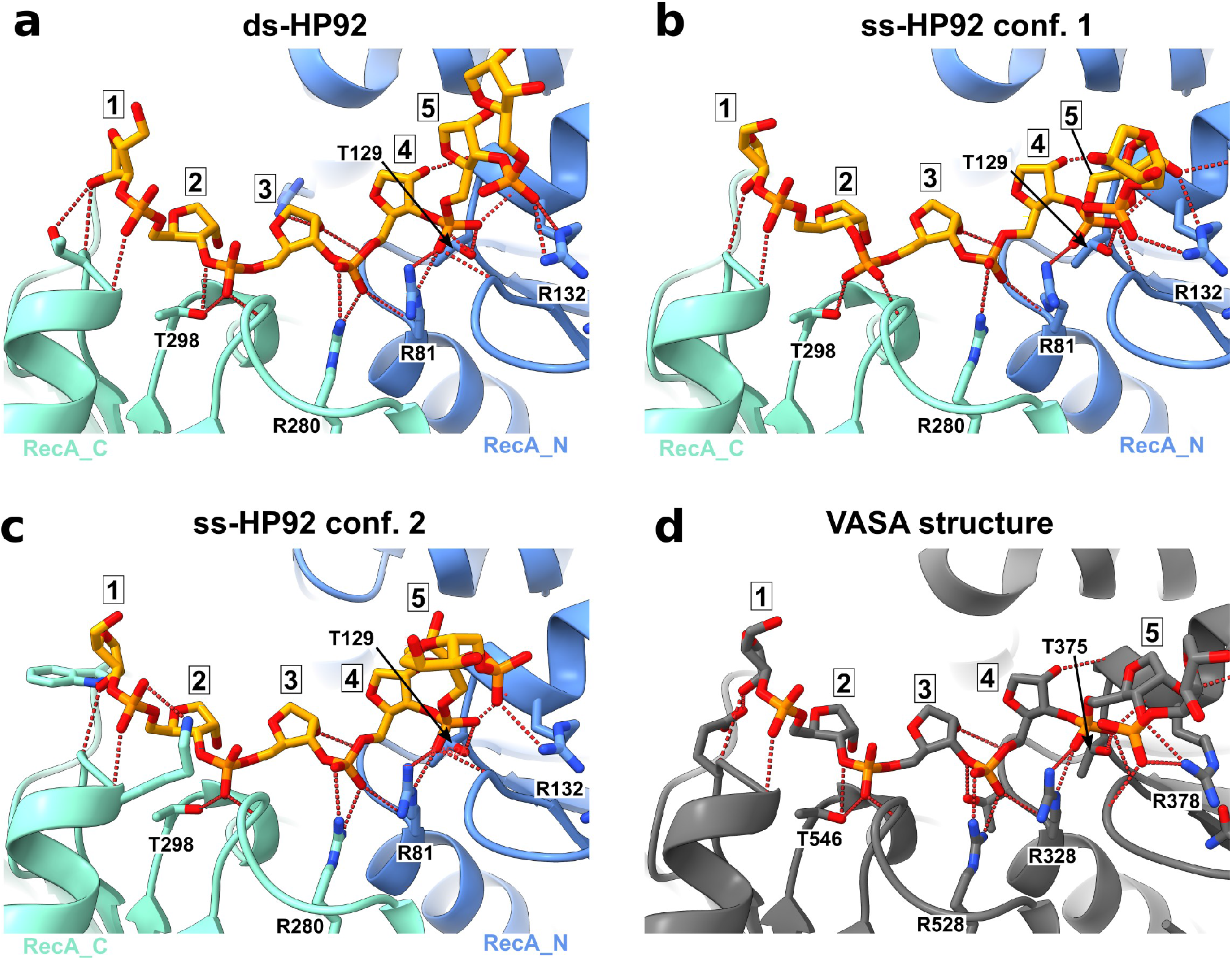
Recognition of the RNA backbone of nt 1-5 for DbpA and VASA. Comparison of the recognition of nt in position 1-5 for the DbpA/ds-HP92 complex (chains B/D) (**a**), DbpA/ss-HP92 complex in conformation 1 (chains L/I) (**b**) and conformation 2 (chains A/C) (**c**) and in the VASA/RNA complex (chains D/H) (**d**). Only the RNA backbone is shown for clarity. Hydrogen bonds between RNA and protein are indicated by by dashed red lines. The nucleotides are numbered 1-5. Protein side chains that contribute important interactions are labeled: R81/R328 are part of the conserved DEAD-box motif Ia (Sengoku et al., 2006), T129/T375 (DbpA/VASA) and R132/R378 (DbpA/VASA) are part of the conserved DEAD-box motif Ib (Sengoku et al., 2006), R280/R528 (DbpA/VASA) are part of the QxxR motif (Sengoku et al., 2006), T546/T298 (DbpA/VASA) are part of motif (Sengoku et al., 2006).

**Figure 7 - figure supplement 1:**
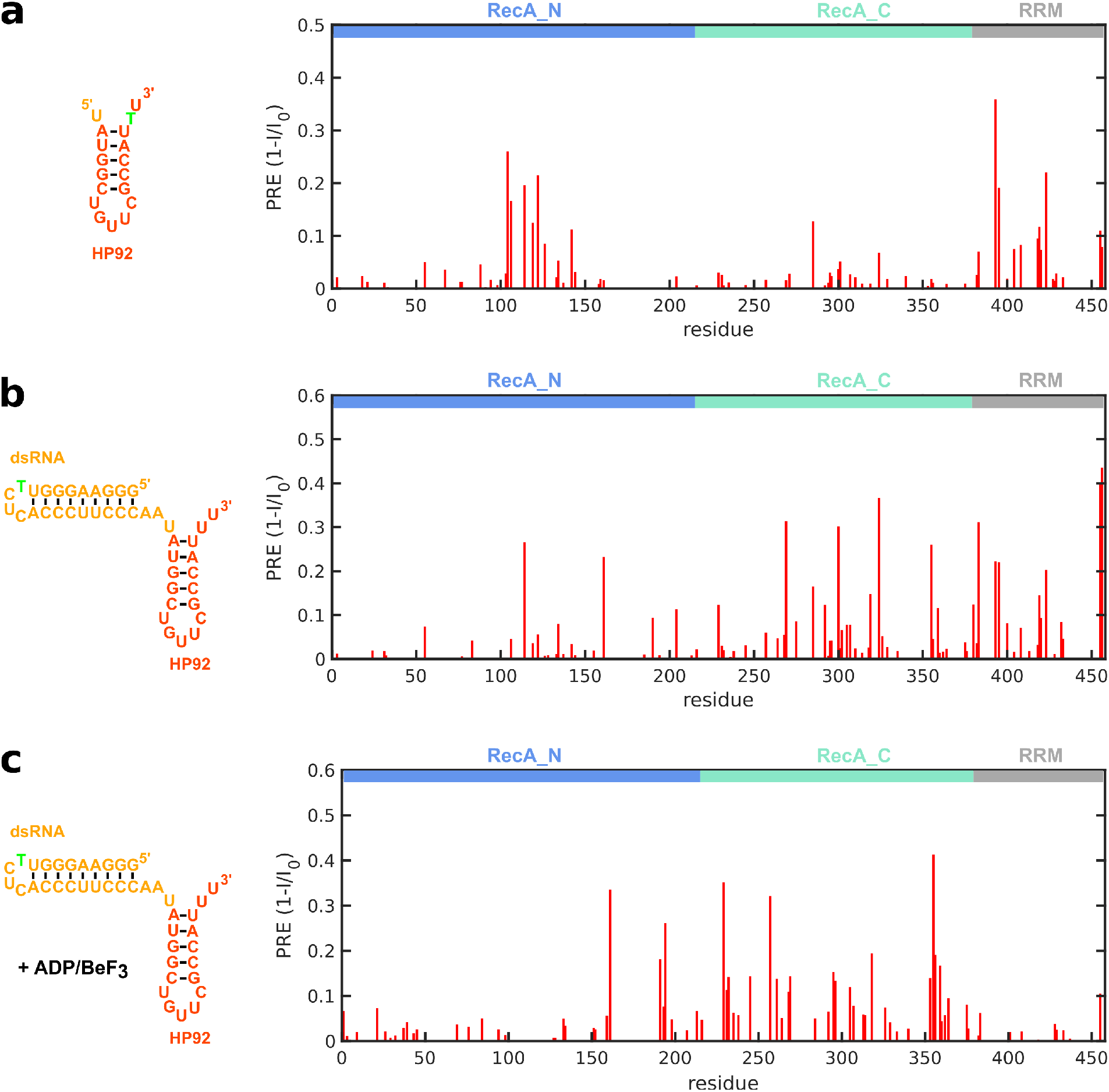
PRE-measurements. (**a**) HP92 RNA construct used for PRE-measurements (left). T (green) indicates the position of the 4-thiouridine residue that is used to couple the nitroxide spin label to the RNA. The determined PRE values (1-I/I_0_) are plotted versus the sequence (right). Increased values indicate the spatial proximity of the spin label. (**b**) hp-HP92 RNA construct used for PRE-measurements (left). T (green) indicates the position of the 4-thiouridine residue that is used to couple the nitroxide spin label to the RNA. The determined PRE values in the absence of ADP/BeF_3_ (1-I/I_0_) are plotted versus the sequence (right). (**c**) hp-HP92 RNA construct used for PRE-measurements (left). T (green) indicates the position of the 4-thiouridine residue that is used to couple the nitroxide spin label to the RNA. The determined PRE values in the presence of ADP/BeF_3_ (1-I/I_0_) are plotted versus the sequence (right).

**Figure 8 - figure supplement 1:**
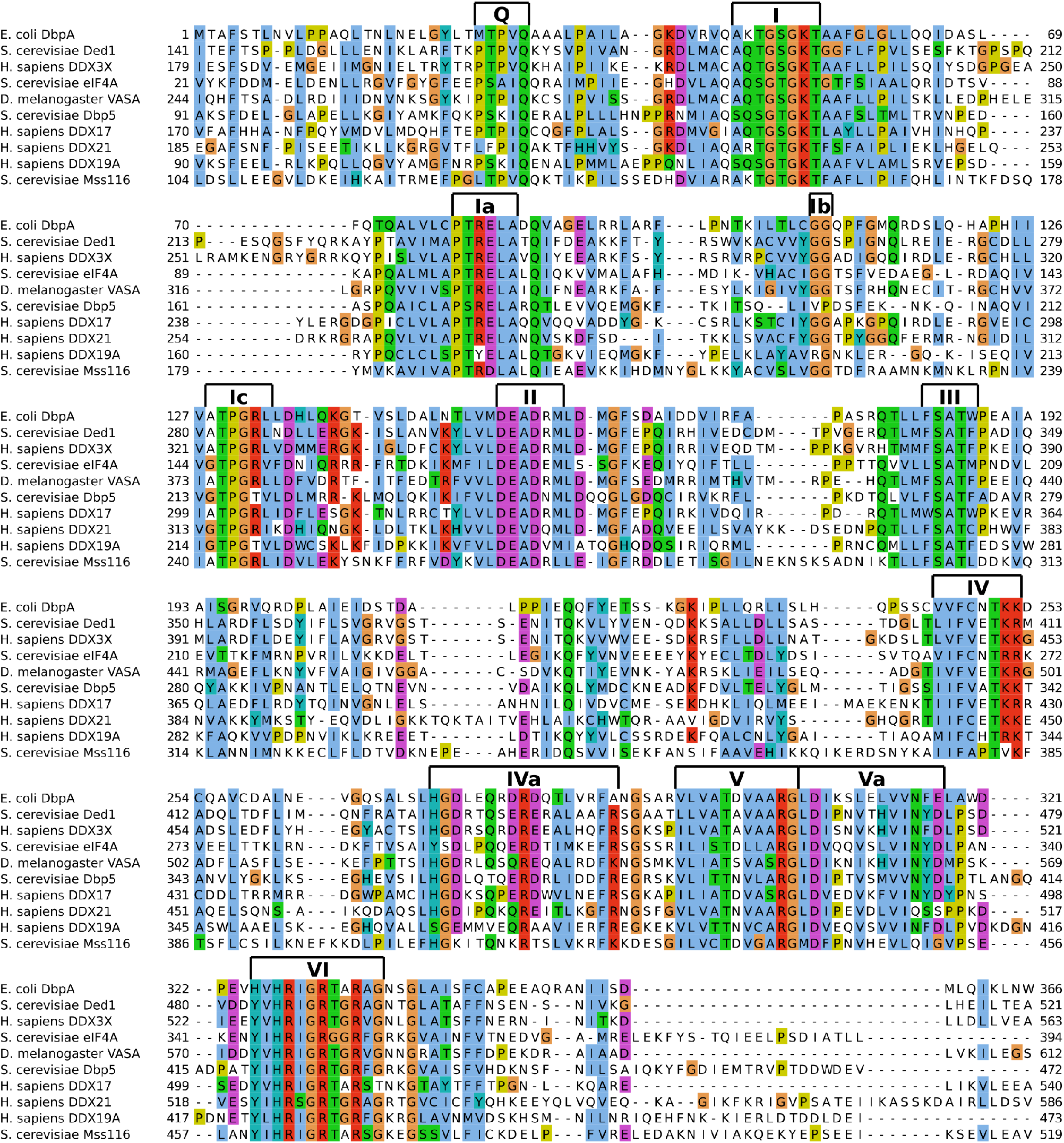
Sequence comparison for DbpA and unrelated DEAD-box helicases. Sequence alignments of the RecA core domains of DbpA (residues 1-366) (top) and nine unrelated DEAD-box helicases. The highly conserved sequence motifs (Q, I-VI (Putnam and Jankowsky, 2013)) of DEAD-box helicases are indicated on top. The sequence alignment was generated using Jalview (Waterhouse et al., 2009) in combination with Clustal Omega (McWilliam et al., 2013).

## Appendix

**Appendix 1 - table 1:**
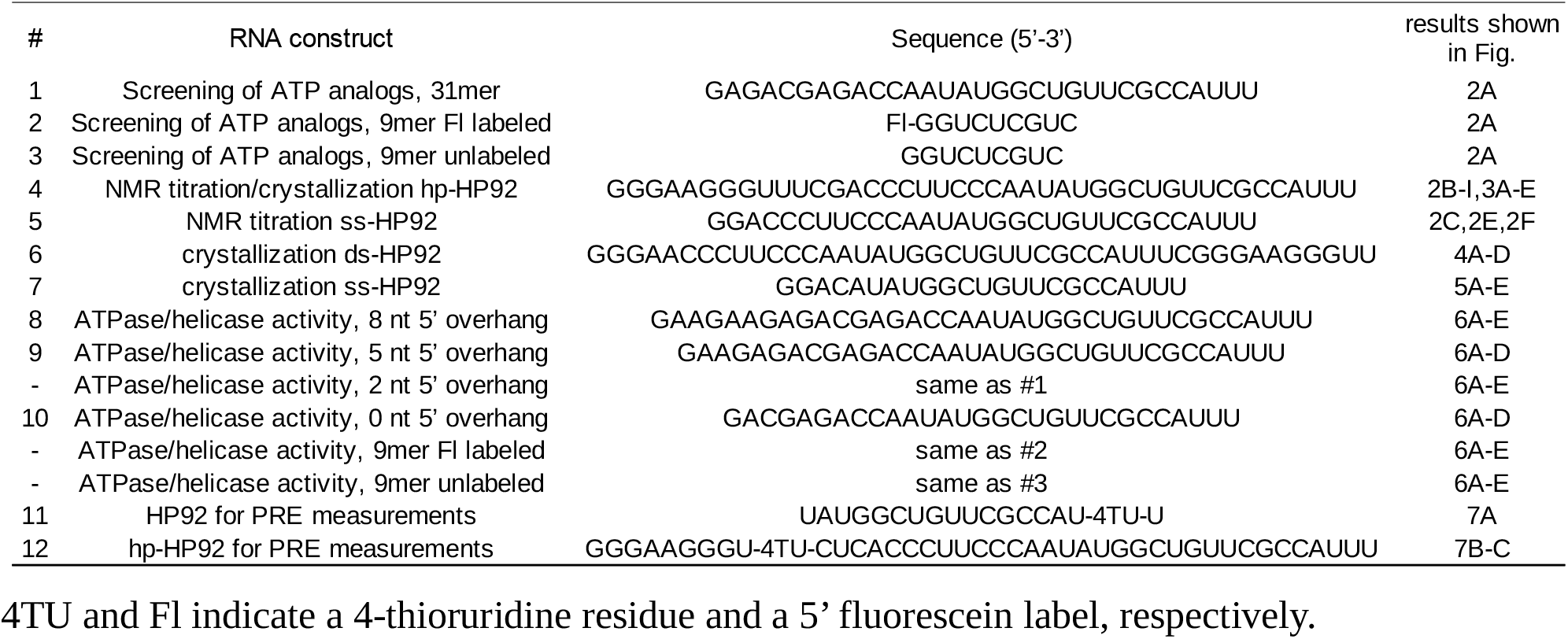
RNA constructs used in this study

## References

Afonine PV, Grosse-Kunstleve RW, Echols N, Headd JJ, Moriarty NW, Mustyakimov M, Terwilliger TC, Urzhumtsev A, Zwart PH, Adams PD. 2012. Towards automated crystallographic structure refinement with phenix.refine. Acta Crystallogr D Biol Crystallogr 68:352–367. doi:10.1107/S0907444912001308

Andreou AZ, Harms U, Klostermeier D. 2019. Single-stranded regions modulate conformational dynamics and ATPase activity of eIF4A to optimize 5’-UTR unwinding. Nucleic Acids Res 47:5260–5275. doi:10.1093/nar/gkz254

Ballut L, Marchadier B, Baguet A, Tomasetto C, Séraphin B, Le Hir H. 2005. The exon junction core complex is locked onto RNA by inhibition of eIF4AIII ATPase activity. Nat Struct Mol Biol 12:861–869. doi:10.1038/nsmb990

Bowers HA, Maroney PA, Fairman ME, Kastner B, Lührmann R, Nilsen TW, Jankowsky E. 2006. Discriminatory RNP remodeling by the DEAD-box protein DED1. RNA 12:903–912. doi:10.1261/rna.2323406

Chen Z, Li Z, Hu X, Xie F, Kuang S, Zhan B, Gao W, Chen X, Gao S, Li Y, Wang Y, Qian F, Ding C, Gan J, Ji C, Xu X-W, Zhou Z, Huang J, He HH, Li J. 2020. Structural Basis of Human Helicase DDX21 in RNA Binding, Unwinding, and Antiviral Signal Activation. Adv Sci (Weinh) 7:2000532. doi:10.1002/advs.202000532

Cianci M, Bourenkov G, Pompidor G, Karpics I, Kallio J, Bento I, Roessle M, Cipriani F, Fiedler S, Schneider TR. 2017. P13, the EMBL macromolecular crystallography beamline at the low- emittance PETRA III ring for high- and low-energy phasing with variable beam focusing. J Synchrotron Radiat 24:323–332. doi:10.1107/S1600577516016465

Clore GM, Tang C, Iwahara J. 2007. Elucidating transient macromolecular interactions using paramagnetic relaxation enhancement. Curr Opin Struct Biol 17:603–616. doi:10.1016/j.sbi.2007.08.013

Collins R, Karlberg T, Lehtiö L, Schütz P, van den Berg S, Dahlgren L-G, Hammarström M, Weigelt J, Schüler H. 2009. The DEXD/H-box RNA helicase DDX19 is regulated by an {alpha}-helical switch. J Biol Chem 284:10296–10300. doi:10.1074/jbc.C900018200

Del Campo M, Lambowitz AM. 2009. Structure of the Yeast DEAD box protein Mss116p reveals two wedges that crimp RNA. Mol Cell 35:598–609. doi:10.1016/j.molcel.2009.07.032

Delaglio F, Grzesiek S, Vuister GeertenW, Zhu G, Pfeifer J, Bax A. 1995. NMRPipe: A multidimensional spectral processing system based on UNIX pipes. Journal of Biomolecular NMR 6. doi:10.1007/BF00197809

Diges CM, Uhlenbeck OC. 2001. Escherichia coli DbpA is an RNA helicase that requires hairpin 92 of 23S rRNA. EMBO J 20:5503–5512. doi:10.1093/emboj/20.19.5503

Emsley P, Cowtan K. 2004. Coot: model-building tools for molecular graphics. Acta Crystallogr D Biol Crystallogr 60:2126–2132. doi:10.1107/S0907444904019158

Fairman-Williams ME, Guenther U-P, Jankowsky E. 2010. SF1 and SF2 helicases: family matters. Curr Opin Struct Biol 20:313–324. doi:10.1016/j.sbi.2010.03.011

Guillerez J, Lopez PJ, Proux F, Launay H, Dreyfus M. 2005. A mutation in T7 RNA polymerase that facilitates promoter clearance. Proc Natl Acad Sci USA 102:5958–5963. doi:10.1073/pnas.0407141102

Güntert P, Mumenthaler C, Wüthrich K. 1997. Torsion angle dynamics for NMR structure calculation with the new program DYANA. J Mol Biol 273:283–298. doi:10.1006/jmbi.1997.1284

Hardin JW, Hu YX, McKay DB. 2010. Structure of the RNA binding domain of a DEAD-box helicase bound to its ribosomal RNA target reveals a novel mode of recognition by an RNA recognition motif. J Mol Biol 402:412–427. doi:10.1016/j.jmb.2010.07.040

Henn A, Cao W, Licciardello N, Heitkamp SE, Hackney DD, De La Cruz EM. 2010. Pathway of ATP utilization and duplex rRNA unwinding by the DEAD-box helicase, DbpA. Proc Natl Acad Sci USA 107:4046–4050. doi:10.1073/pnas.0913081107

Jarmoskaite I, Russell R. 2014. RNA helicase proteins as chaperones and remodelers. Annu Rev Biochem 83:697–725. doi:10.1146/annurev-biochem-060713-035546

Kabsch W. 2010. Integration, scaling, space-group assignment and post-refinement. Acta Crystallogr D Biol Crystallogr 66:133–144. doi:10.1107/S0907444909047374

Keller R. 2004. The computer aided resonance assignment tutorial. Goldau: Cantina Verl.

Kiianitsa K, Solinger JA, Heyer W-D. 2003. NADH-coupled microplate photometric assay for kinetic studies of ATP-hydrolyzing enzymes with low and high specific activities. Anal Biochem 321:266–271. doi:10.1016/s0003-2697(03)00461-5

Linder P, Jankowsky E. 2011. From unwinding to clamping - the DEAD box RNA helicase family. Nat Rev Mol Cell Biol 12:505–516. doi:10.1038/nrm3154

Liu F, Putnam A, Jankowsky E. 2008. ATP hydrolysis is required for DEAD-box protein recycling but not for duplex unwinding. Proc Natl Acad Sci U S A 105:20209–20214. doi:10.1073/pnas.0811115106

Mallam AL, Del Campo M, Gilman B, Sidote DJ, Lambowitz AM. 2012. Structural basis for RNA- duplex recognition and unwinding by the DEAD-box helicase Mss116p. Nature 490:121– 125. doi:10.1038/nature11402

McCoy AJ, Grosse-Kunstleve RW, Adams PD, Winn MD, Storoni LC, Read RJ. 2007. Phaser crystallographic software. J Appl Crystallogr 40:658–674. doi:10.1107/S0021889807021206

Montpetit B, Thomsen ND, Helmke KJ, Seeliger MA, Berger JM, Weis K. 2011. A conserved mechanism of DEAD-box ATPase activation by nucleoporins and InsP6 in mRNA export. Nature 472:238–242. doi:10.1038/nature09862

Mueller M, Wang M, Schulze-Briese C. 2012. Optimal fine ϕ-slicing for single-photon-counting pixel detectors. Acta Crystallogr D Biol Crystallogr 68:42–56. doi:10.1107/S0907444911049833

Ngo TD, Partin AC, Nam Y. 2019. RNA Specificity and Autoregulation of DDX17, a Modulator of MicroRNA Biogenesis. Cell Rep 29:4024–4035.e5. doi:10.1016/j.celrep.2019.11.059

Pettersen EF, Goddard TD, Huang CC, Meng EC, Couch GS, Croll TI, Morris JH, Ferrin TE. 2021. UCSF ChimeraX: Structure visualization for researchers, educators, and developers. Protein Sci 30:70–82. doi:10.1002/pro.3943

Putnam AA, Jankowsky E. 2013. DEAD-box helicases as integrators of RNA, nucleotide and protein binding. Biochim Biophys Acta 1829:884–893. doi:10.1016/j.bbagrm.2013.02.002

Raj S, Bagchi D, Orero JV, Banroques J, Tanner NK, Croquette V. 2019. Mechanistic characterization of the DEAD-box RNA helicase Ded1 from yeast as revealed by a novel technique using single-molecule magnetic tweezers. Nucleic Acids Res 47:3699–3710. doi:10.1093/nar/gkz057

Ren Y, Schmiege P, Blobel G. 2017. Structural and biochemical analyses of the DEAD-box ATPase Sub2 in association with THO or Yra1. Elife 6:e20070. doi:10.7554/eLife.20070

Richardson JS, Schneider B, Murray LW, Kapral GJ, Immormino RM, Headd JJ, Richardson DC, Ham D, Hershkovits E, Williams LD, Keating KS, Pyle AM, Micallef D, Westbrook J, Berman HM, RNA Ontology Consortium. 2008. RNA backbone: consensus all-angle conformers and modular string nomenclature (an RNA Ontology Consortium contribution). RNA 14:465–481. doi:10.1261/rna.657708

Rogers GW, Richter NJ, Merrick WC. 1999. Biochemical and Kinetic Characterization of the RNA Helicase Activity of Eukaryotic Initiation Factor 4A. Journal of Biological Chemistry 274:12236–12244. doi:10.1074/jbc.274.18.12236

Rossi P, Xia Y, Khanra N, Veglia G, Kalodimos CG. 2016. 15N and 13C- SOFAST-HMQC editing enhances 3D-NOESY sensitivity in highly deuterated, selectively [1H,13C]-labeled proteins. J Biomol NMR 66:259–271. doi:10.1007/s10858-016-0074-5

Russell R, Jarmoskaite I, Lambowitz AM. 2013. Toward a molecular understanding of RNA remodeling by DEAD-box proteins. RNA Biol 10:44–55. doi:10.4161/rna.22210

Samatanga B, Klostermeier D. 2014. DEAD-box RNA helicase domains exhibit a continuum between complete functional independence and high thermodynamic coupling in nucleotide and RNA duplex recognition. Nucleic Acids Res 42:10644–10654. doi:10.1093/nar/gku747

Schanda P, Kupce E, Brutscher B. 2005. SOFAST-HMQC experiments for recording two- dimensional heteronuclear correlation spectra of proteins within a few seconds. J Biomol NMR 33:199–211. doi:10.1007/s10858-005-4425-x

Sengoku T, Nureki O, Nakamura A, Kobayashi S, Yokoyama S. 2006. Structural Basis for RNA Unwinding by the DEAD-Box Protein Drosophila Vasa. Cell 125:287–300. doi:10.1016/j.cell.2006.01.054

Sharpe Elles LM, Sykes MT, Williamson JR, Uhlenbeck OC. 2009. A dominant negative mutant of the E. coli RNA helicase DbpA blocks assembly of the 50S ribosomal subunit. Nucleic Acids Res 37:6503–6514. doi:10.1093/nar/gkp711

Snoussi K, Leroy JL. 2001. Imino proton exchange and base-pair kinetics in RNA duplexes. Biochemistry 40:8898–8904. doi:10.1021/bi010385d

Sun Y, Atas E, Lindqvist LM, Sonenberg N, Pelletier J, Meller A. 2014. Single-molecule kinetics of the eukaryotic initiation factor 4AI upon RNA unwinding. Structure 22:941–948. doi:10.1016/j.str.2014.04.014

Theissen B, Karow AR, Köhler J, Gubaev A, Klostermeier D. 2008. Cooperative binding of ATP and RNA induces a closed conformation in a DEAD box RNA helicase. Proc Natl Acad Sci USA 105:548–553. doi:10.1073/pnas.0705488105

Tran EJ, Zhou Y, Corbett AH, Wente SR. 2007. The DEAD-box protein Dbp5 controls mRNA export by triggering specific RNA:protein remodeling events. Mol Cell 28:850–859. doi:10.1016/j.molcel.2007.09.019

Tsu CA, Uhlenbeck OC. 1998. Kinetic analysis of the RNA-dependent adenosinetriphosphatase activity of DbpA, an Escherichia coli DEAD protein specific for 23S ribosomal RNA. Biochemistry 37:16989–16996. doi:10.1021/bi981837y

Wang S, Hu Y, Overgaard MT, Karginov FV, Uhlenbeck OC, McKay DB. 2006. The domain of the Bacillus subtilis DEAD-box helicase YxiN that is responsible for specific binding of 23S rRNA has an RNA recognition motif fold. RNA 12:959–967. doi:10.1261/rna.5906

Wong EV, Cao W, Vörös J, Merchant M, Modis Y, Hackney DD, Montpetit B, De La Cruz EM. 2016. P(I) Release Limits the Intrinsic and RNA-Stimulated ATPase Cycles of DEAD-Box Protein 5 (Dbp5). J Mol Biol 428:492–508. doi:10.1016/j.jmb.2015.12.018

Wurm JP. 2020. Assignment of the Ile, Leu, Val, Met and Ala methyl group resonances of the DEAD-box RNA helicase DbpA from E. coli. Biomol NMR Assign 15:121–128. doi:10.1007/s12104-020-09994-z

Wurm JP, Glowacz K-A, Sprangers R. 2021. Structural basis for the activation of the DEAD-box RNA helicase DbpA by the nascent ribosome. Proc Natl Acad Sci U S A 118:e2105961118. doi:10.1073/pnas.2105961118

Xiol J, Spinelli P, Laussmann MA, Homolka D, Yang Z, Cora E, Couté Y, Conn S, Kadlec J, Sachidanandam R, Kaksonen M, Cusack S, Ephrussi A, Pillai RS. 2014. RNA clamping by Vasa assembles a piRNA amplifier complex on transposon transcripts. Cell 157:1698–1711. doi:10.1016/j.cell.2014.05.018

Yang Q, Del Campo M, Lambowitz AM, Jankowsky E. 2007. DEAD-box proteins unwind duplexes by local strand separation. Mol Cell 28:253–263. doi:10.1016/j.molcel.2007.08.016

